# Loss of TET function in T regulatory cells yields ex-Treg cells biased toward T follicular helper cells, causing autoimmune diseases through autoantibody production

**DOI:** 10.1101/2025.08.29.673187

**Authors:** Kazumasa Suzuki, Leo J. Arteaga-Vazquez, Bruno Villalobos Reveles, Lot Hernández-Espinosa, Isaac F. López-Moyado, Atsushi Onodera, Daniela Samaniego-Castruita, Ferhat Ay, Arlet Lara-Custodio, Patrick G. Hogan, Hugo Sepulveda, Anjana Rao

**Affiliations:** Center for Autoimmunity and Inflammation, La Jolla Institute for Immunology, La Jolla, CA, 92037, USA; Center for Cancer Immunotherapy, La Jolla Institute for Immunology, La Jolla, CA, 92037, USA; Moores Cancer Center, UC San Diego, La Jolla, CA, 92037, USA; Department of Pharmacology, UC San Diego, La Jolla, CA, 92161, USA; Sanford Consortium for Regenerative Medicine, La Jolla, CA, 92037, USA; Institute for Advanced Academic Research (IAAR), Chiba University, Chiba, 263-8522, Japan; Research Institute of Disaster Medicine (RIDM), Chiba University, Chiba, 260-8670, Japan; Center for Human Immunological Diseases and Therapy Development (cCHID), Chiba University, Chiba, 260-8670, Japan; Department of Pediatrics, University of California San Diego, La Jolla, CA, 92093, USA; Program in Immunology, UC San Diego, La Jolla, CA, 92093, USA; Laboratory of Transcription and Epigenetics, Institute of Biomedical Sciences, Universidad Andres Bello, Santiago, Chile

## Abstract

T regulatory cells (Treg cells) express the transcription factor FOXP3 and maintain immune homeostasis by attenuating effector responses. Treg cells are prone to lose FOXP3 and convert to pathological ‘ex-Treg’ cells under conditions of strong or chronic inflammation. One mechanism for loss of FOXP3 expression involves increased DNA methylation of intronic enhancers *CNS1* and *CNS2* in the *Foxp3* locus; these enhancers are maintained in a demethylated state by TET enzymes, 5-methylcytosine (5mC) dioxygenases that generate 5-hydroxymethylcytosine (5hmC) and other oxidized methylcytosines that are essential intermediates in all pathways of DNA demethylation. We previously showed that FOXP3^+^ Treg cells from *Tet2/3*-deficient (*Tet2/3 DKO*) mice displayed increased methylation of *CNS1* and *CNS2* and converted to FOXP3-negative ex-Treg cells considerably more efficiently than WT Treg cells. Here we extend our previous analysis of *Foxp3-Cre Tet2/3^fl/fl^* mice. We classified the mice as DKO- moderate or DKO-severe based on the total number of leukocytes in the spleen and peripheral lymph nodes and investigated the phenotypic and molecular basis for the progressive inflammation occurring in these mice. RNA-seq as well as histological and immunocytochemical analyses showed a striking expansion of T follicular helper (Tfh) cells and plasma cells in *Tet2/3 DKO*-severe mice. RNA-seq analyses also revealed increased induction of interferon-stimulated genes (ISGs) in CD4^+^ FOXP3^-^ T cells from these mice, and single-cell (sc) RNA-seq analyses suggested strongly that this was due to skewed differentiation of both *Tet2/3 DKO* FOXP3^+^ Treg cells and *Tet2/3 DKO* FOXP3^−^ ex-Treg cells into Tfh-like cells. Base-resolution “6-base” sequencing showed the expected loss of 5hmC and increased 5mC in Tfh cells purified from *Tet2/3 DKO*-severe mice, and suggested that the observed bias in gene expression patterns could arise both from a direct increase in methylation of essential enhancers stemming from TET deficiency, or because methylation interfered with binding of methylation-sensitive transcriptional regulators including CTCF.

## INTRODUCTION

T regulatory cells (Treg cells) maintain immune homeostasis and immunological self-tolerance and suppress pathologic inflammation in both humans and mice (1–8). The lineage-determining transcription factor FOXP3 controls the development and suppressive function of Treg cells (9,10). FOXP3-expressing Treg cells are typically stable in steady state conditions *in vivo* (11) but can lose FOXP3 and convert to pathological cells called ‘ex-Treg’ cells in strong or chronic inflammatory environments (11–20). Ex-Treg cells lose the ability to suppress inflammation and can additionally convert to effector CD4^+^ T cells of different lineages – Th1, Th2, Th17 and T follicular helper (Tfh) cells that secrete pro-inflammatory cytokines (IFN-γ, IL-4, IL-17, IL-21) – thereby promoting chronic inflammation and the development of autoimmune/inflammatory diseases (11–20). Ex-Treg cells have been suggested to promote autoimmune reactions in several mouse models of disease, including type 1 diabetes (14), graft-versus-host disease (16), multiple sclerosis (17), and autoimmune arthritis (18,19). But it is still unclear how ex-Treg cells acquire inflammatory functions after loss of FOXP3.

The stability of FOXP3 expression during mouse Treg cell differentiation is regulated by two conserved noncoding sequence elements, *CNS1* and *CNS2*, located in the first intron of the *Foxp3* gene (5’ of the first coding exon) (21–23). *CNS2*, also termed Treg specific demethylated region (TSDR) (24), controls the stability of *Foxp3* expression in a manner linked to the DNA modification status of *CNS2* (24–27). CpG sites in the *Foxp3 CNS2* element are predominantly unmethylated (C/5fC/5caC) in Treg cells, but fully methylated (5mC/5hmC) in naive T cells (21,24,27–32). The TET dioxygenases TET2 and TET3 act redundantly to deposit 5-hydroxymethylcytosine (5hmC) at the *CNS1* and *CNS2* enhancers during Treg cell differentiation, resulting in their demethylation (loss of 5-methylcytosine, 5mC) (28,33). Combined disruption of the *Tet2* and *Tet3* genes in developing thymocytes of *CD4-Cre Tet2/3^floxed/floxed^* (*Tet2/3^flfl^*) mice resulted in striking antigen-dependent expansion of iNKT cells, together with massive inflammation due to partial loss of T regulatory cells in the thymus and their almost complete loss in the periphery (34). The loss of Treg cells in TET-deficient mice was at least partly due to reduced Treg stability due to loss of FOXP3 expression, which in turn was due to increased DNA methylation of *CNS1* and *CNS2* (28,30,33,35). Addition of the TET activator vitamin C during *in vitro* differentiation of mouse and human iTreg cells increased TET enzymatic activity, maintained the *CNS1* and *CNS2* enhancers in a demethylated state, and increased the stability of FOXP3 expression (28,31). Importantly, both *CNS2*-deficient and *Tet2/3*-deficient Treg cells lose FOXP3 in a manner that depends on cell division (21,22,28). The loss of FOXP3 can be ameliorated by treatment with IL-2 or certain other cytokines (22) or the combination of IL-2, retinoic acid and Vitamin C (28,36).

We previously showed that combined TET2/TET3 deficiency in Treg cells from *Foxp3-Cre Tet2/3^flfl^* mice led to the progressive development of an autoimmune/ inflammatory disease that was fatal by 8–22 weeks of age (33,35). Treg cells from these mice displayed increased methylation of the intronic enhancers *CNS1* and *CNS2* in the *Foxp3* gene, albeit to a lesser extent than observed in *CD4-Cre Tet2/3^flfl^* mice, presumably reflecting the fact that CD4 is expressed earlier than FOXP3 during Treg development. In this early study, we analyzed the molecular phenotypes of only two 14-week-old *Foxp3-Cre Tet2/3^flf^* mice, and RNA-seq data from CD4^+^ FOXP3^-^ T cells from one of these two mice showed upregulation of Tfh-related and Th17-related genes (33). Moreover, Treg cells from *Foxp3^Cre^ Tet2/3^flfl^* mice were more prone to lose FOXP3 expression and become ‘ex-Treg’ cells (33). Notably, when we transferred total CD4^+^ T cells from *Foxp3-Cre Tet2/3^flfl^* mice into recipient immunocompetent mice (33), we found that these cells were uniformly able to elicit a dominant inflammatory disease, despite the presence of wildtype Treg cells in the recipient mice. These data emphasized two points: first, that TET proteins are essential for maintenance of Treg cell stability and immune homeostasis in mice, and second, that the inflammatory phenotype in the transfer experiments was driven, at least in part, by ex-Treg cells that acquired effector function (33).

In this study, we expanded our analysis of the role of TET dioxygenases in Treg and ex-Treg cells from *Foxp3-Cre Tet2/3^fl/fl^* mice. Phenotypically, the mice showed clear evidence of splenomegaly and lymphadenopathy that we classified as moderate or severe based on the total number of leukocytes in the spleen. The frequencies and numbers of total CD4^+^ T cells and CD4^+^ FOXP3^+^ Treg cells increased strikingly in DKO-severe compared to WT mice. As the disease advanced from DKO-moderate to DKO-severe, the mice showed increased T cell activation as judged by a progressive decrease in the numbers of naïve CD4^+^ T cells, and increased inflammation based on increased expression of interferon-stimulated genes (ISGs). Histological and immunocytochemical analyses, together with flow cytometry, showed increased lymphocyte infiltration into many organs, increased numbers of neutrophils and plasma cells in DKO-severe mice, and expansion of cytotoxic CD8^+^ T cells with increased expression of perforin and granzyme B. By performing bulk and single-cell (sc) RNA-seq on CD4^+^ T cells from WT and DKO-severe mice, we found increased expression of Tfh-associated genes, which was confirmed by flow cytometry revealing increased expression of Tfh surface markers both in FOXP3^+^ Treg cells and FOXP3^-^ ex-Treg cells of *Foxp3Cre DKO* mice. Changes in 5mC and 5hmC distribution presented a complex picture depending on the genes involved, suggesting that increased methylation due to TET deficiency could reflect direct methylation of essential enhancers or decreased binding of transcriptional repressors such as CTCF. We conclude that FOXP3-negative ex-Treg cells from mice with selective deficiency in *Tet2* and *Tet3* in Treg cells show biased conversion to Tfh cells and cause systemic inflammation with variable onset whose severity progressively increases with age.

## RESULTS

### Defining the level of systemic inflammation in *Foxp3-Cre Tet2/3^fl/fl^* mice

At 14 weeks of age, *Foxp3-Cre Tet2/3^fl/fl^* mice could be divided into two groups distinguished by moderate or severe splenomegaly and lymphadenopathy (**Fig. 1A**), defined as DKO-moderate and DKO-severe respectively depending on whether they had fewer than or more than 4 x 10^8^ splenocytes and peripheral lymphocytes (from cervical and inguinal) (**Fig. 1B**). The DKO-severe group contained a higher proportion of females (9/14, 64%) compared to the DKO-moderate group (3/11, 27%) (**Fig. 1C**), suggesting that females are more likely to develop severe symptoms in *Foxp3-Cre Tet2/3^fl/fl^*mice.

**Fig. 1.**
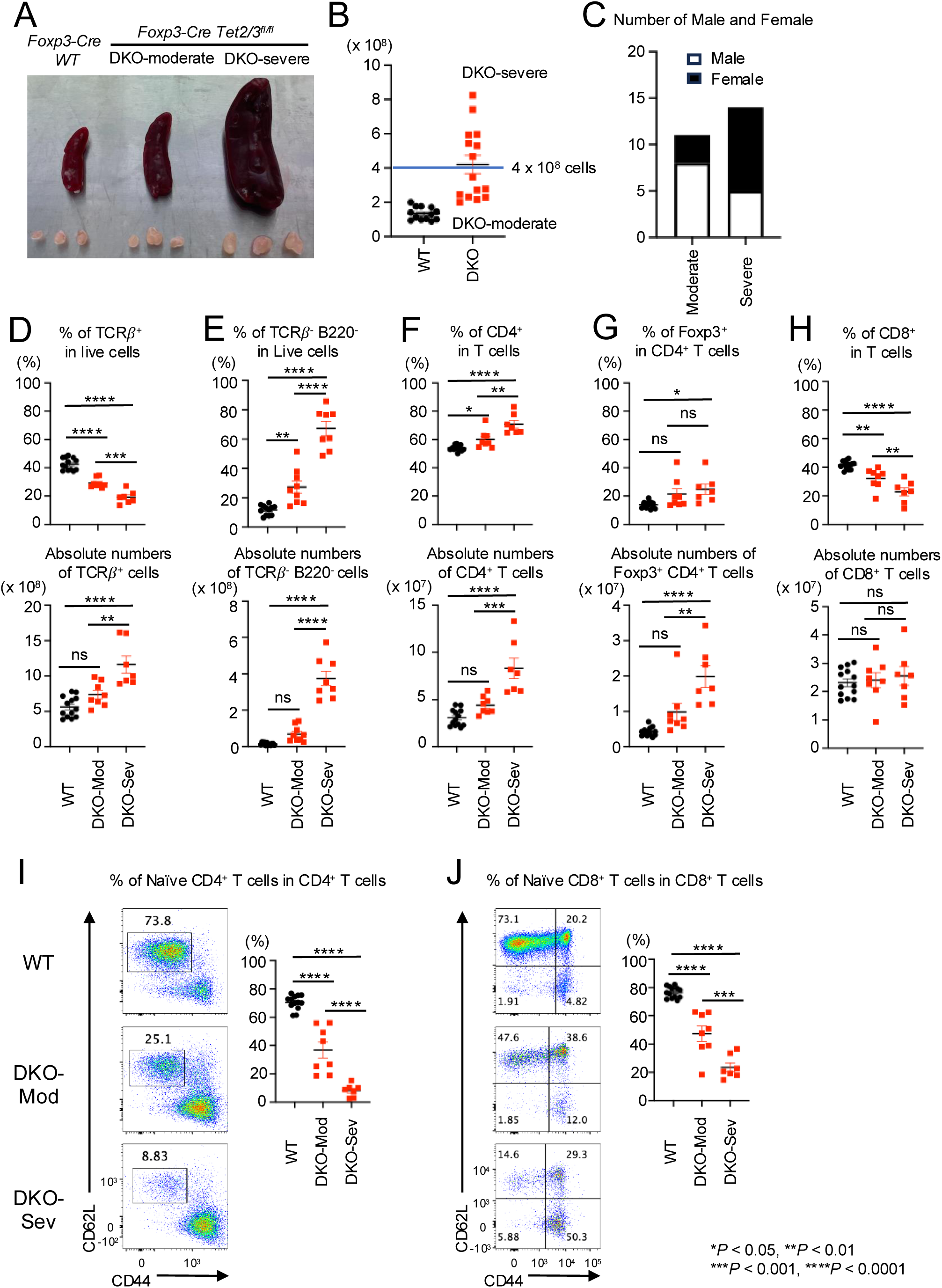
DKO-severe *Foxp3-Cre Tet2/3 ^fl/fl^* mice display CD4^+^ cells and TCR𝛽^-^ B220^-^ cells proliferation with strong inflammation. **A.** Representative pictures of spleen and peripheral lymph nodes (cervical and inguinal) from *Foxp3Cre WT*, DKO- moderate and DKO-severe *Foxp3-Cre Tet2/3^fl/fl^* mice (14-weeks-old). **B.** Absolute cell number of pooled spleen and peripheral lymph nodes (cervical and inguinal) of *Foxp3Cre WT*, DKO-moderate and DKO-severe *Foxp3-Cre Tet2/3^fl/fl^*mice (14-weeks-old). **C.** Number of male and female in DKO-moderate and DKO-severe in *Foxp3-Cre Tet2/3^fl/fl^* mice (14-weeks-old) **D-H.** Quantification of the frequency (*Top*) and absolute number (*bottom*) in pooled spleen and peripheral lymph nodes (cervical and inguinal) from *Foxp3Cre WT*, *Foxp3-Cre Tet2/3^fl/fl^*mice (14-weeks-old). **D.** Frequency of TCRβ^+^ cells in live cells and absolute number. **E.** Frequency of TCRβ^-^ B220^-^ cells in live cells and absolute number. **F.** Frequency of CD4^+^ cells in TCRβ^+^ cells and absolute number. **G.** Frequency of YFP(FOXP3)^+^ cells in TCRβ^+^ CD4^+^ cells and absolute number. **H.** Frequency of CD8^+^ cells in TCRβ^+^ cells and absolute number. **I-J.** Representative flow cytometry plots of CD62L^high^ CD44^low^ cells in TCRβ^+^ CD4^+^ cells (**I**) or TCRβ^+^ CD8^+^ cells (**J**) and quantification of the frequency of the cells in pooled spleen and peripheral lymph nodes (cervical and inguinal) from *Foxp3Cre WT*, *Foxp3-Cre Tet2/3^fl/fl^* mice (14-weeks-old). Error bars show mean ± SEM. from at least three independent experiments. Statistical analysis was performed using one-way ANOVA (\**P* < 0.05, \*\**P* <0.01, \*\*\**P* < 0.001, \*\*\*\**P* < 0.0001).

Although the percentage of T cells (TCRβ^+^ cells) among live cells in spleen and lymph nodes decreased, the absolute number of T cells increased in DKO-severe compared to WT or DKO-moderate mice (**Fig. 1D**). This reflected an increase in the frequencies and absolute numbers of cells other than T or B cells (TCRβ^-^ B220^-^ cells) in both DKO- moderate and DKO-severe mice (**Fig. 1E**; also see **Fig. 4A** below). The expansion of CD4^+^ T cells (both frequencies and absolute numbers) showed a slight increase in the DKO-moderate group and a considerable increase in the DKO- severe group compared to WT (**Fig. 1F**). The absolute numbers of *Tet2/3DKO* Treg cells were also increased, especially in DKO-severe mice (**Fig. 1G**); in light of the dominant inflammatory phenotype of *Foxp3Cre Tet2/3^fl/fl^* mice, this finding supports our prior demonstration that Tregs in these mice have largely lost suppressive function, as measured by their decreased ability, compared to wildtype Tregs, to correct the inflammation induced by *scurfy* cells in T cell and bone marrow adoptive transfer experiments (33). The absolute numbers of CD8^+^ cells (as a fraction of all T cells) did not change in spleen and lymph nodes of DKO-moderate or DKO-severe compared to WT mice (**Fig. 1H**); however, the percentage of naïve cells (CD62L^high^ CD44^lo^) was dramatically decreased in both CD4^+^ and CD8^+^ T cell populations in *Foxp3-Cre Tet2/3^fl/fl^* mice, especially DKO-severe mice, indicating that the majority of the T cells were activated or had become memory cells (**Fig. 1I, J**).

### Apparent skewing of *Tet2/3 DKO* CD4^+^ YFP(FOXP3)^-^ ex-Treg cells towards Tfh- and Th17-like cells

We performed RNA sequencing on CD4^+^ YFP(FOXP3)^-^ T cells from WT, DKO-moderate or DKO-severe mice. WT CD4^+^ YFP(FOXP)^-^ T cells, which contained ∼70% naïve T cells (**Fig. 1I**, *top*), clustered relatively closely (less than 9% variance in a PCA plot, **Suppl. Fig. 1A**), whereas CD4^+^ YFP(FOXP)^-^ T cells from either DKO-moderate or DKO-severe *Foxp3-Cre Tet2/3^fl/fl^*mice showed considerably more variability – likely because they are a heterogenous cell population containing *Tet2/3DKO* ex-Treg cells arising from loss of FOXP3 stability as well as conventional (non-Treg) CD4^+^ T cells mostly displaying an activated or memory phenotype (**Fig. 1I**, *middle and bottom panels*). The DKO- moderate and DKO-severe CD4^+^ T cells showed a significant upregulation of Tfh-related genes (*Bcl6*, *Cxcr5*, *Pdcd1*, *Tox2*, *Maf, Batf, Ascl2, Il21*, *Il4*, *Klrb1c, Nfkb1*), Th17-related genes (*Rorc*, *Il17a*, *Il17f*) and cell cycle genes (*Cdkn1a*, *Cdkn2a*, *E2f7*, *E2f8 (data not shown*) compared to WT (MA plots, **Suppl. Fig. 1B, C**), suggesting that a proportion of *Tet2/3DKO* Treg cells had converted into proliferating ex-Treg cells with Tfh-like or Th17-like phenotypes.

When differentially expressed genes were extracted and clustered into a heatmap (**Fig. 2A**), most Tfh-related (*Bcl6*, *Cxcr5*, *Pdcd1*, *Tox2*, *Maf, Batf, Il21, Nfkb1*) and Th17-related (*Rorc*, *Il17a*, *Il17f*) genes were expressed more strongly in DKO-severe than in DKO-moderate or WT CD4^+^ T cells (heatmap cluster 3). Moreover, *Foxo1* – a suppressor of Tfh differentiation – was less highly expressed in DKO cells compared to WT (heatmap cluster 1). These results suggested that loss of FOXP3 expression in *Tet2/3 DKO* ex-Treg cells might result in expansion of ex-Treg cells with gene expression patterns resembling those of Tfh or Th17 cells.

**Fig. 2.**
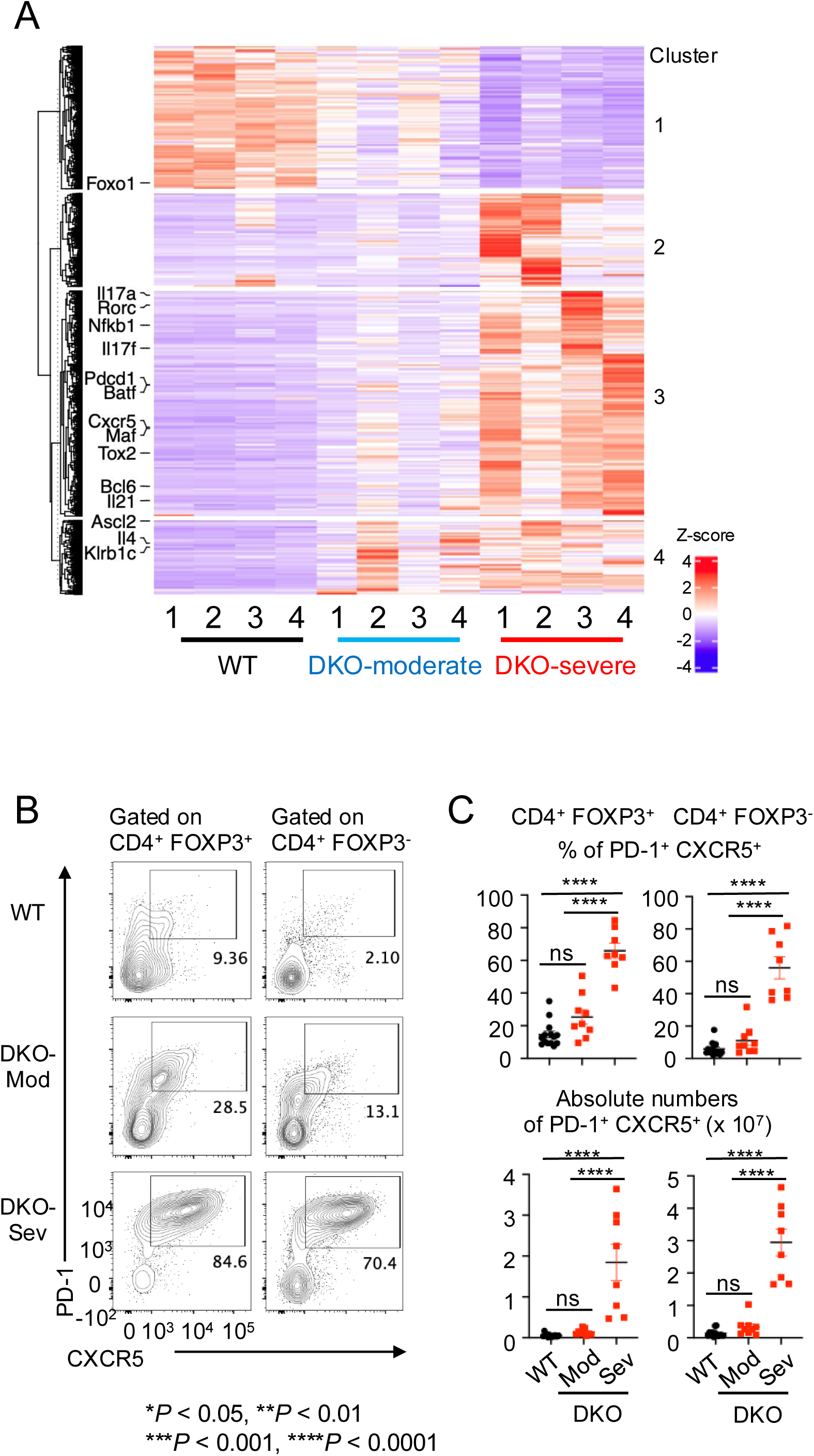
Tfh-like cells expanded in in DKO-severe *Foxp3-Cre Tet2/3 ^fl/fl^* mice in bulk RNA sequencing and flow cytometry analysis. **A.** Heatmap for the expression of clustered differential expressed genes (DEGs). CD4^+^ YFP(FOXP3)^-^ T cells were sorted from pooled spleen and peripheral lymph nodes (cervical and inguinal) from *Foxp3Cre WT* mice and *Foxp3- Cre Tet2/3^fl/fl^* mice (14-weeks-old) and RNA sequencing was performed (n=4). DEGs were extracted and clustered. Heatmap for the expression of clustered DEGs was generated. Tfh related (*Bcl6*, *Cxcr5*, *Pdcd1*, *Tox2*, *Maf, Batf, Nfkb1, Ascl2, Il21, Il4, Klrb1C, Foxo1*), Th17 related (*Rorc, Il17a, Il17f*) genes were indicated. **B.** Flow cytometry analysis of PD-1^+^ CXCR5^+^ cells (gated on TCRβ^+^ CD4^+^ YFP(FOXP3)^+^ or TCRβ^+^ CD4^+^ YFP(FOXP3)^-^ cells) in pooled spleen and peripheral lymph nodes (cervical and inguinal) from 13-15 weeks old *Foxp3Cre WT* and *Foxp3-Cre Tet2/3^fl/fl^* mice. **C.** Quantification of the frequency of PD-1^+^ CXCR5^+^ cells in TCRβ^+^ CD4^+^ YFP(FOXP3)^+^ or TCRβ^+^ CD4^+^ YFP(FOXP3)^-^ cells and absolute number in pooled spleen and peripheral lymph nodes (cervical and inguinal). Error bars show mean ± SEM. from at least three independent experiments. Statistical analysis was performed using one-way ANOVA (\**P* < 0.05, \*\**P* <0.01, \*\*\**P* < 0.001, \*\*\*\**P* < 0.0001).

We confirmed the striking and progressive expansion of Tfh-like PD-1^+^ CXCR5^+^ cells in DKO-severe mice by flow cytometry of both residual CD4^+^ YFP(FOXP3)^+^ Treg cells and CD4^+^ YFP(FOXP3)**^−^**cells (containing expanded ex-Treg cells). A remarkable 55-65% of both residual CD4^+^ FOXP3^+^ and CD4^+^ FOXP3^-^ cells in DKO-severe mice were PD-1^+^ CXCR5^+^ Tfh-like cells (range 45-85%, **Fig. 2B, C**). Consistent with this finding, there was a remarkable increase in the frequency of Fas^+^ GL-7^+^ GC B cells in B220^+^ cells in spleens of DKO-severe compared to DKO-moderate or WT mice (**Suppl. Fig. 2J**), even though the frequencies of total B220^+^ cells declined (**Suppl. Fig. 2I**).

In contrast, a relatively small and variable proportion of cells in total splenic CD4^+^ T cells of DKO-severe mice produced IL-17A upon PMA/ionomycin stimulation (1-12%, **Suppl. Fig. 2A**, *left*, **Suppl. Fig. 2D**). The frequencies of CD8^+^ T cells that showed increased expression of perforin and granzyme B, and that produced IFN-γ upon PMA/ionomycin stimulation, increased in DKO-severe compared to WT mice, suggesting expansion of bystander cytotoxic CD8^+^ T cells (**Suppl. Fig. 2B, C, F-H**).

Taken together, these results indicate a strong expansion of Tfh-like cells and GC B cells in *Tet2/3* deficient mice that is especially evident in mice with a DKO-severe phenotype. Some of these cells may arise directly from bystander CD4^+^ T cells, whereas others may arise from expanded ex-Treg cells that have lost FOXP3 and have acquired skewed gene expression patterns that resemble those of PD-1^+^ CXCR5^+^ Tfh-like cells.

### Greater expansion of Tfh-like cells compared to Th17-like cells in DKO-severe *Foxp3-Cre Tet2/3 ^fl/fl^* mice

To address this question further, we compared patterns of gene expression at the single-cell level in WT and DKO- severe TCRβ^+^ T cells (**Fig. 3**). Single-cell (sc) RNA-sequencing distinguished 19 clusters in UMAP analysis (**Fig. 3A**, explained in **Suppl Fig. 3**). *Cd4^+^* and *Cd8^+^*cells occupy the *left* and *right* arms of the UMAP plot respectively (**Fig. 3B**). Cells in clusters 0, 5 were primarily WT naïve CD4^+^, cells in clusters 1, 2 were primarily WT naïve CD8^+^ T cells, and cells in clusters 6 (mostly DKO) and 9 (mostly WT) were Treg cells based on expression of *Cd4* and *Foxp3* (**Fig. 3A, Suppl Fig. 3**). Interestingly, *Tet2/3 DKO Foxp3*^+^ Treg cells (Cluster 6) expressed not only *Foxp3* and other Treg signature genes (*Il2ra*, *Nrp1*, *Ctla4, Ikzf4*), but also Tfh-related genes (*Bcl6*, *Cxcr5*, *Pdcd1*, *Icos*, *Tox2*, *Maf, Batf*), suggesting that conversion to Tfh-like ex-Treg cells was initiated in *Tet2/3DKO Foxp3*^+^ Treg cells prior to loss of FOXP3 due to reduced transcription and expression stability (**Fig. 3C**). This finding was consistent with our previous study showing that Tfh-related genes were upregulated in Treg cells compared to WT Treg cells in a mouse with a DKO- severe phenotype of two *Tet2/3DKO* (*Foxp3Cre*) mice analyzed; the other mouse was DKO-moderate as defined in Fig. 1B (33). This conclusion was further supported by analysis of cluster 3, which contained primarily DKO-severe, Foxp3^-^ ex-Treg cells: almost all cells in this cluster expressed Tfh-related genes very strongly (**Fig. 3C**).

**Fig. 3.**
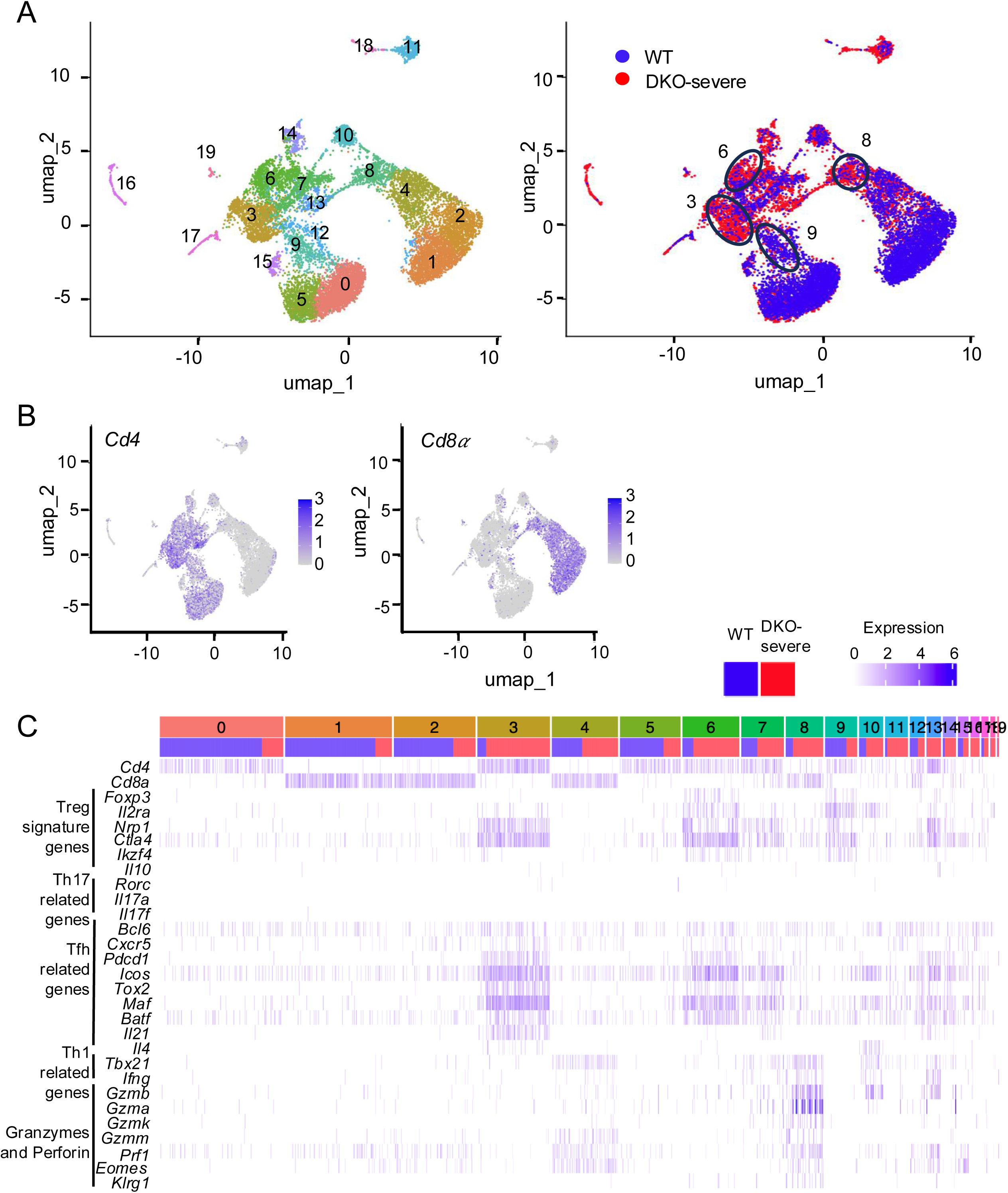
Tfh-like cells expanded in in DKO-severe *Foxp3-Cre Tet2/3 ^fl/fl^* mice at the single-cell level analysis. **A-C.** Single-cell (sc) RNA sequencing for T cells from WT or DKO-severe mice. TCRβ^+^ cells were sorted from pooled spleen and peripheral lymph nodes (cervical and inguinal) from *Foxp3Cre WT* mice and *DKO-severe Foxp3-Cre Tet2/3^fl/fl^* mice (14-weeks-old) and sc-RNA sequencing was performed (n=1). **A, B.** The UMAP plot showed 0-19 clusters (**A**, *left*), distribution of WT (Blue) and *Tet2/3DKO* (Red) population (**A**, *right*) and expression of *Cd4* (**B**, *left*), *Cd8α* (**B**, *right*). **C.** Expression map of indicated genes. Blue column: Cells from WT mice, Red column: Cells from DKO-severe mice.

To determine whether Tfh cells in *Foxp3-Cre Tet2/3^fl/fl^*mice were derived from *Tet2/3*-deficient ex-Treg cells, we measured *Tet2* expression in Treg and Tfh cells from WT and *Foxp3-Cre Tet2/3^fl/fl^* mice by qPCR. Relative to WT Treg cells, *Tet2/3 DKO* Treg cells did not detectably express *Tet2*, whereas *Tet2^+/-^* Treg cells expressed about half the amount of *Tet2* compared to WT Treg cells (**Suppl. Fig. 1D**, *left*). Tfh cells from DKO-moderate and DKO-severe *Foxp3-Cre Tet2/3^fl/fl^* mice expressed significantly lower and undetectable levels of *Tet2* respectively compared to other CD4^+^ cell populations in these mice, including WT non-Treg cells, non-Tfh cells not expressing Tfh surface markers and naïve CD4^+^ cells (**Suppl. Fig. 1D**, *right*). *Tet2* expression was particularly reduced in Tfh cells from DKO-severe mice (**Suppl. Fig. 1D**, *right*), suggesting that almost all Tfh cells in DKO-severe *Foxp3-Cre Tet2/3^fl/fl^* mice were derived from *Tet2/3DKO* ex-Treg cells.

*Rorc* and *Il17a/Il17f* expression were barely detected in sc-RNA seq of *Tet2/3 DKO* cells (**Fig. 3C**), although they were clearly detected by bulk RNA-seq of the entire DKO-severe cell population (**Fig. 2A, Suppl. Fig. 1C**). In an attempt to resolve this discrepancy, we performed flow cytometry on stimulated CD4^+^ and CD8^+^ T cells from WT, DKO- moderate and DKO-severe mice. The average percentage of IL-17A-secreting CD4^+^ T cells in spleen and peripheral lymph nodes of DKO-severe *Foxp3-Cre Tet2/3^fl/fl^* mice after PMA-ionomycin stimulation was only ∼5%, although this represented a clear increase above the percentages observed in WT or DKO-moderate mice (1 and 2% respectively; **Suppl. Fig. 2A**, *left*, **Suppl. Fig. 2D**). These data indicated that Th17 cells were not a major population in *Tet2/3 DKO* ex-Treg cells. It has been reported that ex-Treg cells possess Th1 characteristics (12,14,17), but the percentages of IFN-γ^+^ cells (presumably Th1 cells) were decreased rather than increased in CD4^+^ cells from *Tet2/3 DKO*-severe mice (**Suppl. Fig. 2A**, *right*, **Suppl. Fig. 2E**). In contrast, however, CD8^+^ T cells did not expand (**Fig. 1H**) but showed increased cytotoxic function in *Tet2/3 DKO*-severe compared to WT or DKO-moderate mice, as judged both by sc- RNA-seq for *Tbx21*, *Ifng*, *Granzymes* (*Gzma*, *Gzmb*, *Gzmk, Gzmm*) and *Prf1* encoding perforin (**Fig. 3C**, cluster 8 in sc-RNA-seq expression map) and by flow cytometry for IFN-γ, granzyme B and perforin (**Suppl. Fig. 2B, C, F-H**). IL- 21 produced by Tfh cells has been shown to induce differentiation of CD8^+^ T cells into cytotoxic effector cells by upregulating key molecules such as IFN-γ and granzyme B in anti-tumor immunity or viral infection (37,38), suggesting that Tfh-like *Tet2/3 DKO* ex-Treg cells induced cytotoxic CD8^+^ T cell differentiation via IL-21 in *Tet2/3 DKO*-severe mice.

### Expansion of plasma cells and Tfh cells in secondary lymphoid tissues of DKO-severe *Foxp3-Cre Tet2/3 ^fl/fl^* mice

As mentioned above, although the percentage of T cells (TCRβ^+^ cells) in live cells decreased, the absolute number of T cells increased in DKO-severe *Foxp3Cre Tet2/3 ^fl/fl^* mice (**Fig. 1D**). Moreover, the percentage of B220^+^ cells in live cells declined but the absolute number did not change in DKO-severe compared to WT or DKO-moderate mice (**Suppl. Fig. 2I**). The most expanded cell population in spleens and peripheral lymph nodes of DKO-severe were a TCRβ^-^ B220^-^ population (∼70%) (**Fig. 4A**), which contained CD11b^+^ Gr-1^+^ neutrophils (**Fig. 4B** *left***, C**) as well as an expanded population of CD98^+^ CD138^+^ plasma cells (**Fig. 4B** *right*, **Fig. 4D**). Plasma cells showed clear increases in both frequencies and numbers in DKO-severe compared to WT or DKO-moderate mice (**Fig. 4D**). We attribute plasma cell expansion in *Foxp3Cre Tet2/3^fl/fl^* mice to the Tfh activity of the expanded *Tet2/3 DKO* ex-Treg cells, which would be expected to promote B cell activation in germinal center and plasma cell differentiation.

**Fig. 4.**
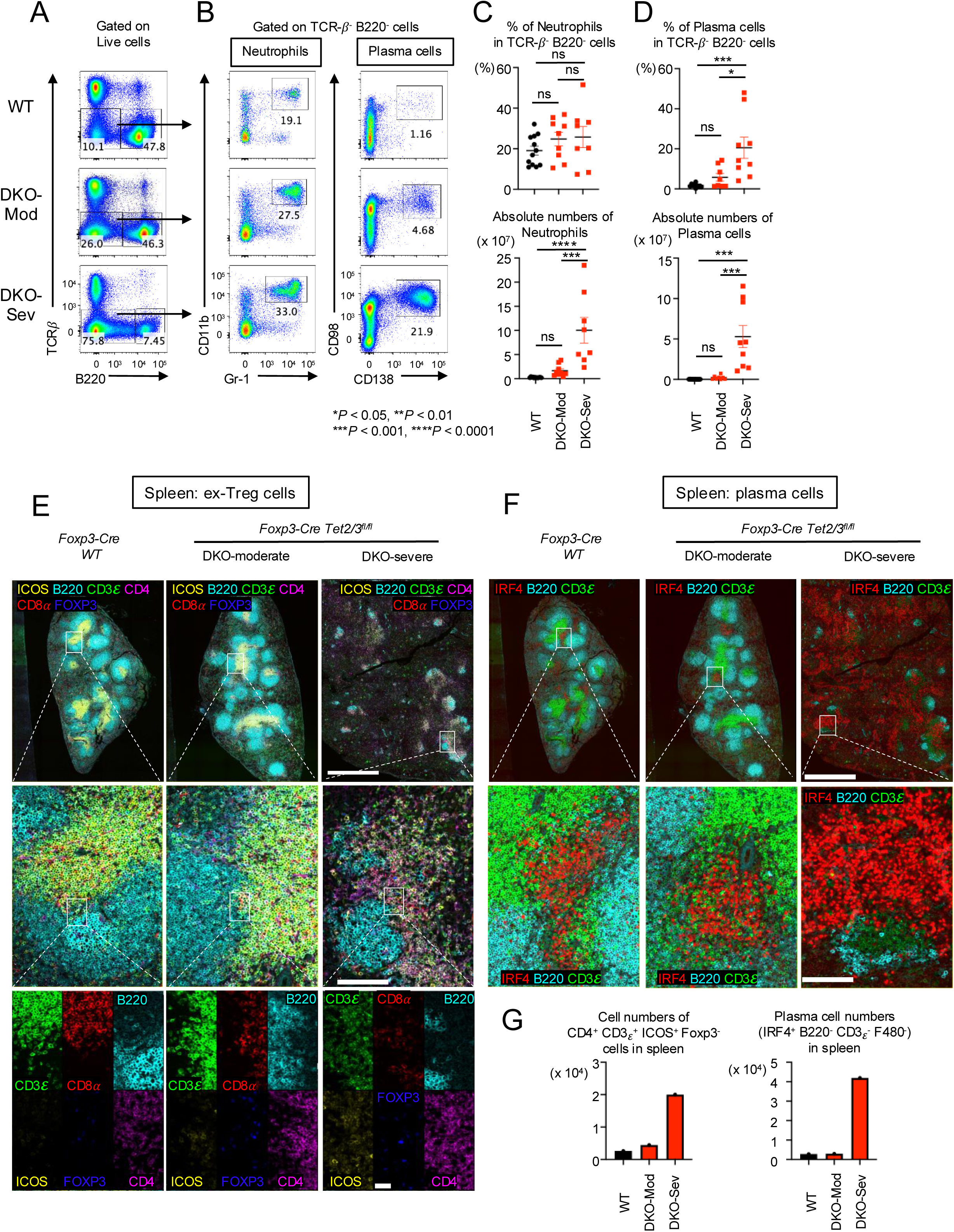
Along with proliferation of Tfh-like *Tet2/3 DKO* ex-Treg cells, plasma cells were diffusely expanded in spleen from DKO-severe mice. **A.** Flow cytometry analysis of TCRβ^-^ B220^-^ cells (gated on live cells) in pooled spleen and peripheral LNs (cervical and inguinal) of from 13-15 weeks old *Foxp3Cre WT* and *Foxp3-Cre Tet2/3^fl/fl^* mice. **B.** Flow cytometry analysis of neutrophils (CD11b^+^ Gr-1^+^ cells) or plasma cells (CD98^+^ CD138^+^ cells) (gated on TCRβ^-^ B220^-^ cells) in pooled spleen and peripheral LNs (cervical and inguinal) of from 13-15 weeks old *Foxp3Cre WT* and *Foxp3-Cre Tet2/3^fl/fl^* mice. **C.** Quantification of the frequency of CD11b^+^ Gr-1^+^ cells in TCRβ^-^ B220^-^ cells and absolute number. **D.** Quantification of the frequency of CD98^+^ CD138^+^ cells in TCRβ^-^ B220^-^ cells and absolute number. Error bars show mean ± SEM. from at least three independent experiments. Statistical analysis was performed using one-way ANOVA (\**P* < 0.05, \*\**P* <0.01, \*\*\**P* < 0.001, \*\*\*\**P* < 0.0001). **E.** Immunostaining of CD3ε, CD8α, B220, ICOS, FOXP3 and CD4 in spleen from 14-week-old *Foxp3Cre WT* and *Foxp3-Cre Tet2/3^fl/fl^* mice. Scale bar; top: 1mm, middle: 100μm, bottom: 25μm. **F.** Immunostaining of IRF4, B220, CD3ε in spleen from 14-week-old *Foxp3Cre WT* and *Foxp3-Cre Tet2/3^fl/fl^* mice. Scale bar; top: 1mm, bottom: 100μm. **G.** The graphs show the cell numbers of CD4^+^ CD3ε^+^ ICOS^+^ FOXP3^-^ cells or IRF4^+^ B220^-^ CD3ε^-^ F480^-^ cells counted in the stained sections from spleen.

We performed immunostaining analysis for Tfh-like *Tet2/3 DKO* ex-Treg cells and plasma cells. In **Fig. 4E**, there are clear T cell and B cell regions (as well as germinal centers (*data not shown)*) in spleens from WT or DKO-moderate mice, but these structures were destroyed in DKO-severe mice (**Fig. 4E**, *top*). In T cells of WT, the ratio of CD4^+^ T cells to CD8^+^ did not differ perceptibly and there were almost no ICOS^+^ cells in WT spleens (**Fig. 4E**, *left middle and bottom panels*); in contrast, the proportion of CD4^+^ cells and ICOS^+^ cells increased slightly in DKO-moderate and significantly in DKO-severe spleen (**Fig. 4E**, *right two panels, middle and bottom*) in the T cell-B cell border region. ICOS is a well-established marker for Tfh cells; given that a substantial number of *Tet2/3 DKO* Treg cells in DKO- severe mice have not yet lost *Foxp3* but nevertheless express *Icos* (clusters 6 in **Fig. 3C**), a likely possibility is that the ICOS^+^ CD4^+^ FOXP3**^-^** cells in *Foxp3Cre Tet2/3 ^fl/fl^* mice are derived from TET-deficient Treg cells that converted into Tfh-like ex-Treg cells that gain ICOS expression (see Discussion).

In Fig. **4F**, we used IRF4 as a plasma cell marker. IRF4^+^ cell clusters were observed in WT and DKO-moderate spleen, but their number and extent were limited (**Fig. 4F**, *left panels*). In contrast, IRF4-positive cells were observed extensively throughout the spleen of DKO-severe mice (**Fig. 4F**, *right panels*), consistent with the increase in frequencies and absolute numbers of CD98^+^ CD138^+^ plasma cells in the spleens of these mice (**4B** *right*, **Fig. 4D**). The absolute numbers of FOXP3^-^ Tfh-like cells and plasma cells in the spleen were also strikingly increased in *Tet2/3 DKO*-severe phenotype mice compared to WT mice (**Fig. 4G**).

Finally, we tested the polyreactivity of serum isolated from WT (n=8), DKO-moderate (n=3) and DKO-severe (n=4) by autoantibody array assay and observed elevated levels of autoantibodies related to autoimmune diseases (**Suppl. Fig. 4**). The assay between groups was conducted using the R package ‘limma’ and multiple comparisons correction was performed (adjusted p value < 0.05 between tested groups). Consistent with plasma cell expansion, the levels of many autoantibodies were elevated variably but substantially in DKO-severe compared to WT or DKO-moderate mice (**Suppl. Fig. 4**).

### Strong inflammation and autoimmune reactions in several tissues of DKO-severe *Foxp3-Cre Tet2/3 ^fl/fl^* mice

We performed multi-tissue necroscopy analysis for WT, DKO-moderate and DKO-severe mice. We evaluated the degree of inflammation using our Inflammation score (0–5) for H&E staining samples from pancreas, colon, skeletal muscle, heart, liver, lung, salivary gland, brain, knee joint, stomach. We observed strong inflammation and lymphocyte infiltration in kidney, liver, lung and pancreas in DKO-severe mice (**Fig. 5A, Suppl. Fig. 5**). In kidneys of WT and DKO- moderate mice, infiltrates were absent or limited to sparse lymphocytes and macrophages. In contrast, DKO-severe mice exhibited prominent mixed cortical interstitial infiltrates enriched in plasma cells, particularly surrounding renal veins near the fornix. In the livers of DKO-severe mice, large numbers of lymphocytes, plasma cells, and macrophages expanded periportal regions and filled hepatic sinusoids. In the lungs, DKO-moderate mice had perivascular and peribronchiolar mixed infiltrates dominated by lymphocytes, consistent with infiltrated inducible BALT (Bronchus-associated lymphoid tissue). These infiltrates were more extensive in DKO-severe mice, forming dense cuffs with increased plasma cell content. In the pancreas, DKO-moderate mice occasionally exhibited lymphocytic infiltration of islets without statistical significance. Pancreata from DKO-severe mice were densely infiltrated by plasma cells, lymphocytes, and macrophages. In the most affected cases, this immune infiltrate completely disrupted normal pancreatic architecture (**Suppl. Fig. 5**).

**Fig. 5.**
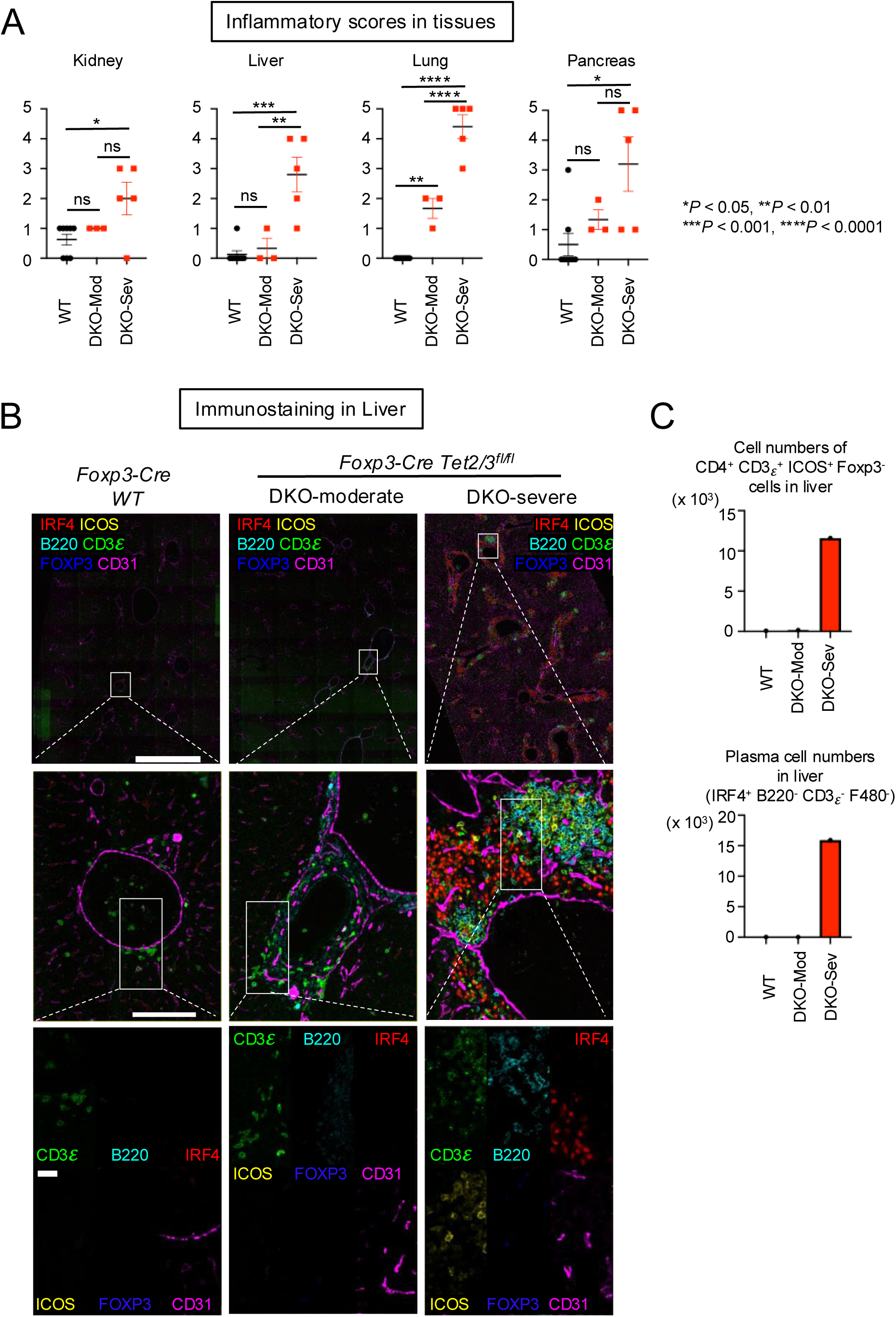
Systemic inflammation accompanied by infiltration of Tfh-like ex-Treg cells and plasma cells damaged organs in DKO-severe mice. **A.** Inflammation scores (0-5) measured from H&E staining. Error bars show mean ± SEM. from at least three independent experiments. Statistical analysis was performed using one-way ANOVA (\**P* < 0.05, \*\**P* <0.01, \*\*\**P* < 0.001, \*\*\*\**P* < 0.0001). **B.** Immunostaining of CD3ε, B220, IRF4, ICOS, FOXP3 and CD31 in liver from 14-week-old *Foxp3Cre WT* and *Foxp3-Cre Tet2/3^fl/fl^* mice. Scale bar; top: 1mm, middle: 100μm, bottom: 25μm.**C.** The graphs show the cell numbers of CD4^+^ CD3ε^+^ ICOS^+^ Foxp3^-^ cells or IRF4^+^ B220^-^ CD3ε^-^ F480^-^ cells counted in the stained sections from liver.

To determine what cell types infiltrated perivascular regions in livers from DKO-severe mice, we performed immunostaining of liver sections. Livers of WT mice showed infiltration of a few CD3ε^+^ cells and DKO-moderate livers showed infiltration of CD3ε^+^ cells but the numbers were not many (**Fig. 5B**, *left two panels*). CD3ε^+^ T cells in DKO- moderate livers did not express ICOS, suggesting that these T cells were not Tfh-like cells (**Fig. 5B**, *middle panels*; **Fig. 5C**, *top*). Moreover, in perivascular regions in livers from DKO-severe mice, CD3ε^+^ ICOS^+^ cells, B220^+^ cells, and IRF4^+^ cells were observed (**Fig. 5B**, *right bottom panel*; **Fig. 5C**, *bottom*). Only a small fraction of *Tet2/3 DKO* CD3ε^+^ ICOS^+^ Tfh-like cells expressed FOXP3 and IRF4^+^ cells were plasma cells whose proliferation was observed in the spleen. Together these results suggested that *Tet2/3 DKO* Tfh-like ex-Treg cells and plasma cells proliferated in secondary lymphoid tissues and migrated to peripheral tissues, possibly exacerbating inflammation directly.

### Upregulation of Interferon stimulated genes (ISGs) in CD4^+^ YFP(FOXP3)^-^ cells from DKO-severe mice

As mentioned above, CD4^+^ YFP(FOXP3)^-^ T cells isolated from spleens of WT mice were primarily (∼70%) naïve CD4+ T cells (**Fig. 1I**, *top panel*), whereas CD4^+^ YFP(FOXP3)**^-^** T cells from DKO-moderate and DKO-severe mice included ∼15% and ∼60% of PD-1^+^ CXCR5^+^ Tfh-like cells respectively (**Fig. 2B, C**). A substantial fraction of which likely corresponded to *Tet2/3 DKO* ex-Treg cells (**Suppl. Fig. 1D**, *right*). To determine whether these cells were responding to an inflammatory environment, we analyzed their expression of interferon-stimulated genes (ISGs) by RNA-seq of sorted CD4^+^ YFP(FOXP3)^-^ T cells.

Indeed, expression of many ISGs was altered in CD4^+^ YFP(FOXP3)**^-^** T cells from DKO-severe and DKO-moderate mice compared to WT mice (**Fig. 6A-C**). The total number of ISGs whose expression was significantly altered, either by upregulation or downregulation, increased substantially from DKO-moderate to DKO-severe (**Fig. 6A, B**), and mRNAs encoding a large number of chemokines, cytokines and their ligands or receptors were upregulated in cells from DKO-moderate mice and to an even greater extent in cells from DKO-severe mice (**Fig. 6C**, *clusters 2-4*). The variability among the DKO mice reflects the variable nature of disease progression in vivo, as also seen in human patients with autoimmune disease.

**Fig. 6.**
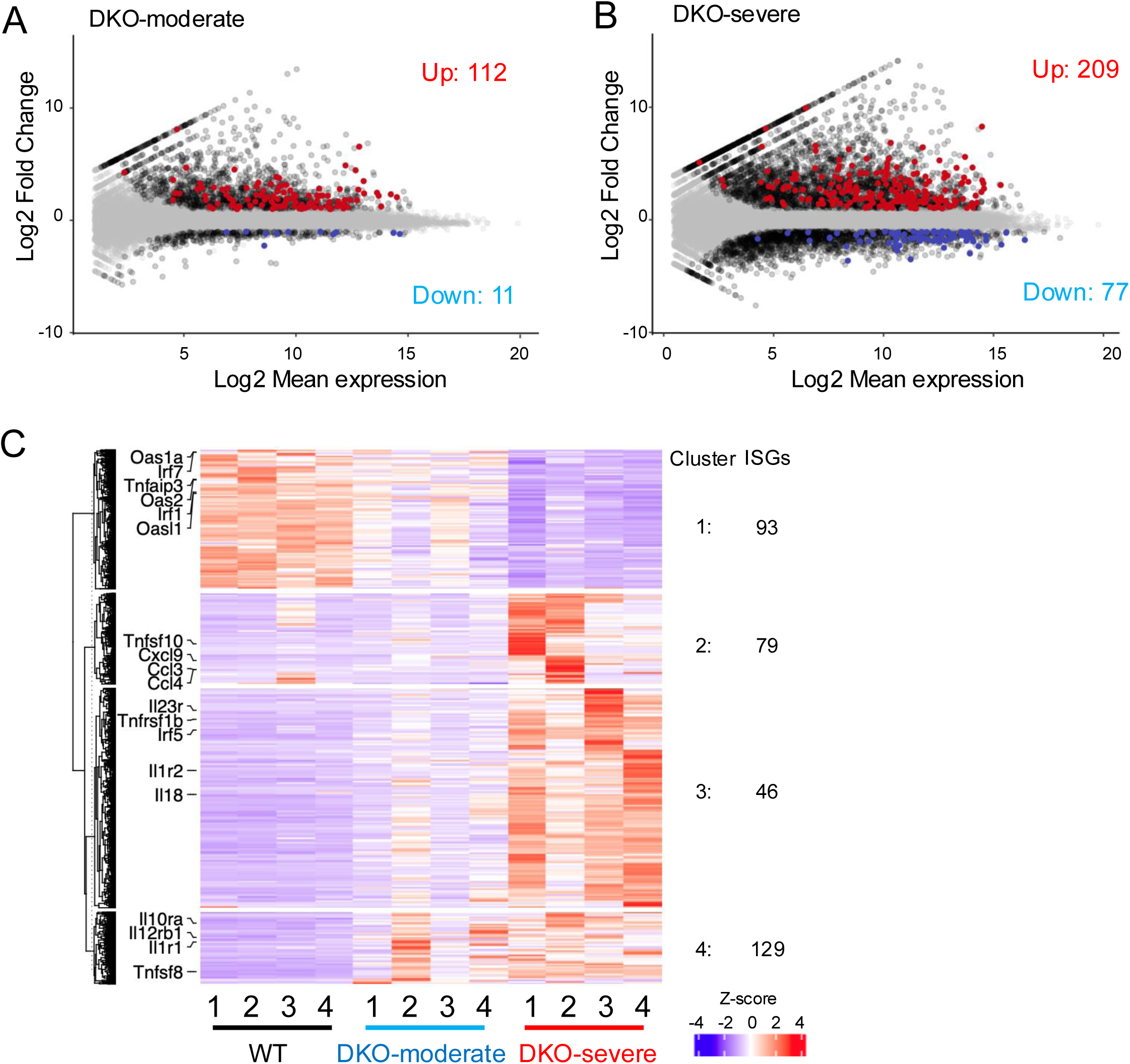
Interferon stimulated genes (ISGs) were up-regulated in CD4^+^ YFP(FOXP3)^-^ T cells from DKO-severe mice. **A-B.** Mean average (MA) plot of differentially expressed ISGs in DKO-moderate (**A**) or DKO-severe (**B**) relative to their expression in WT. ISGs are listed in the “ISGs list (1032 genes)”. RNA-seq analysis was done for CD4^+^ YFP(FOXP3)^-^ T cells from *Foxp3Cre WT* mice and *Foxp3-Cre Tet2/3^fl/fl^* mice (14-weeks-old). **A.** MA-plot of DKO- moderate vs WT. **B.** MA-plot of DKO-severe vs WT. Red: up-regulated ISGs, Blue; down-regulated ISGs, Black; DEGs (no ISGs), Grey: no DEGs. **C**. Heatmap for the expression of clustered differential expressed genes (DEGs). Indicated genes are listed in the “selected ISGs list (44 genes)”.

Notably, *Irf7* and several members of the OAS (2’-5’ oligoadenylate synthase family) were downregulated in CD4^+^ FOXP3**^-^** cells from DKO-moderate and DKO-severe mice compared to WT mice (**Fig. 6C**, *cluster 1*). Of these, OAS1, OAS2, and OAS3 enzymes initiate the degradation of ds-RNA viruses by generating 2’-5’ oligoadenylate which then activates RNase L to degrade the viral ds-RNAs (39), while OASL1 is known to inhibit translation of *Irf7* mRNA (40). Similarly, the *Tnfaip3* gene encodes the TNF-induced protein A20 (TNFAIP3), a remarkable enzyme that inhibits multiple facets of signaling to NFκB that combines binding to K63-linked and linear ubiquitin chains with deubiquitinase activity on K63- and K48-linked ubiquitin chains within a single protein (41). A20 expression is normally induced by TNF, a cytokine expressed at high levels under inflammatory conditions; complete deletion of the *Tnfaip3* gene in mice leads to perinatal lethality from multiorgan inflammation and cachexia, while haploinsufficiency of A20/ TNFAIP3 in humans has been linked to a wide variety of inflammatory conditions (42). Thus, these enzymes are all part of the bona fide anti-viral response, whereas the genes indicated in clusters 2-4, which encode secreted proteins and their ligands and receptors, dictate T cell responses to viral infections.

### *Bcl6* upregulation with concomitant hypermethylation of the *Bcl6* locus in *Tet2/3DKO* Tfh cells

TET enzymes deposit 5hmC at enhancers to maintain them in an active, demethylated state (43,44). We examined DNA cytosine modification patterns in naïve CD4^+^ T cells (CD4^+^ YFP(FOXP3)^-^ CD62L^high^ CD44^low^) from WT and *Tet2/3 DKO* Tfh-like ex-Treg (CD4^+^ YFP(FOXP3)^-^ PD-1^+^ CXCR5^+^) cells of *Tet2/3 DKO*-severe mice (both 14-week-old) using 6-base sequencing, a method that provides information on the four canonical bases A, C, G, T and the modified bases 5mC and 5hmC in a single sequencing run (45) (**Fig. 7**, **Suppl. Fig. 6**). The results revealed several interesting but diverse ways in which methylation changes correlate with, and presumably regulate, changes in gene expression.

**Fig. 7.**
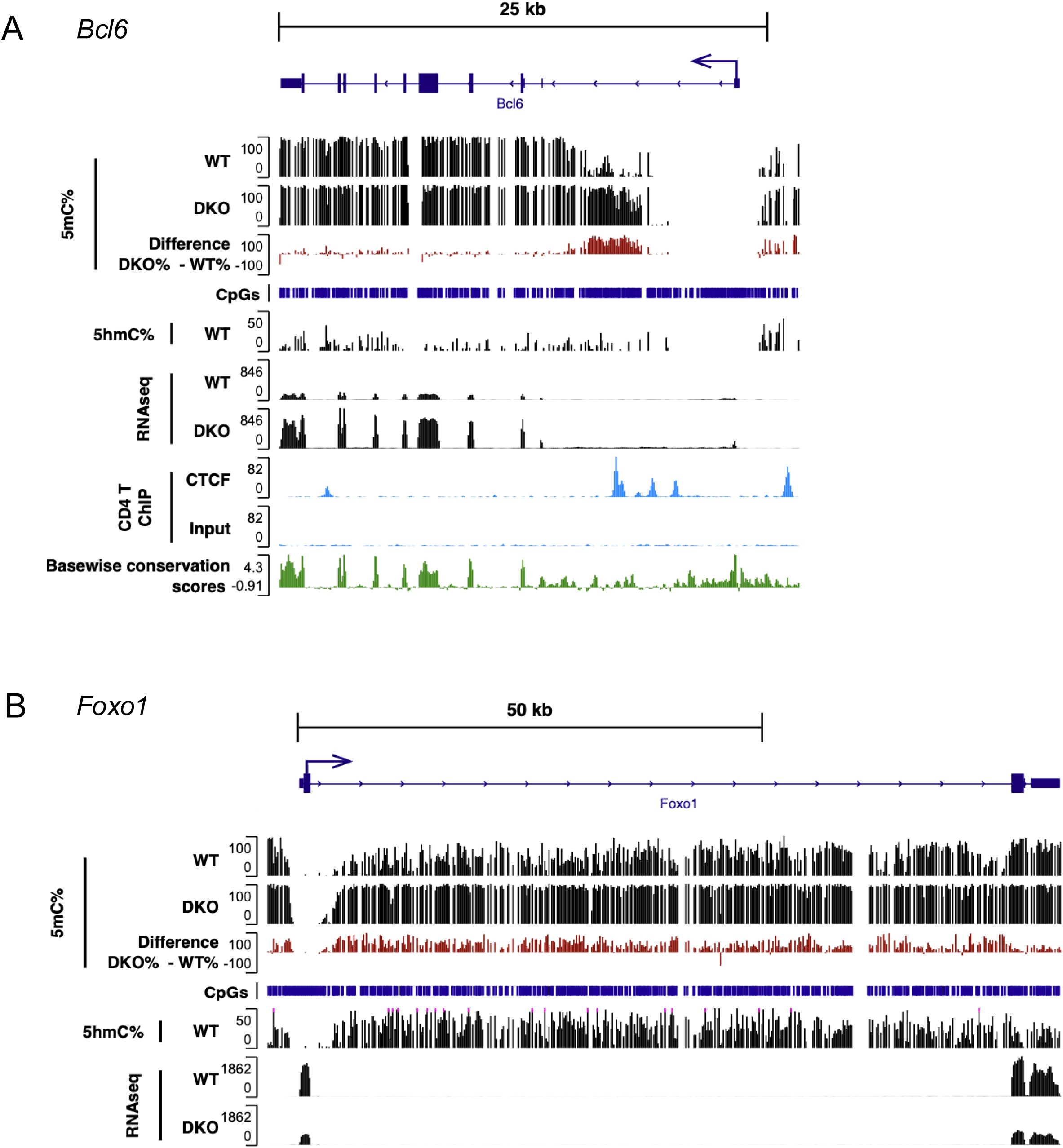
Characteristic hypermethylation accumulated in the first intron of the *Bcl6* locus in *Tet2/3DKO* Tfh cells. **A.** Genome browser views showing 5mC% (Track 1), 5hmC% (Track 2) from 6 base sequencing, gene expression (RNA-seq, track 3), CTCF binding (ChIP-seq, track 4) with CpGs, Basewise conservation scores in *Bcl6* locus. **B.** Genome browser views showing 5mC% (Track 1), 5hmC% (Track 2) from 6 base sequencing, gene expression (RNA- seq, track 3) with CpGs in *Foxo1* locus. 6 base sequencing; WT: naïve CD4^+^ T cells (CD4^+^ YFP(FOXP3)^-^ CD62L^high^ CD44^low^) from *Foxp3Cre WT* mice, DKO: Tfh like cells (CD4^+^ YFP(FOXP3)^-^ PD-1^+^ CXCR5^+^) from DKO-severe *Foxp3- Cre Tet2/3^fl/fl^* mice. RNA seq; WT: CD4^+^ YFP(FOXP3)^-^ T cells from *Foxp3Cre WT* mice, DKO: CD4^+^ YFP(FOXP3)^-^ T cells from DKO-severe *Foxp3-Cre Tet2/3^fl/fl^* mice. ChIP-seq; conventional CD4^+^ T cells were isolated from WT mice and ChIP-seq for CTCF was performed (56).

BCL6 is well-established as a lineage-defining transcription factor for Tfh cells (46–52). We observed a highly significant increase in *Bcl6* mRNA expression in Tfh-like ex-Treg cells in DKO-severe compared to WT naïve CD4^+^ cells (**Suppl. Fig. 1C, Fig. 2A**, **Fig. 3C**), that correlated with a striking increase in methylation of a region in the first intron of the *Bcl6* gene (**Fig. 7A**). This finding is consistent with a previous publication (53,54) showing that CD4^+^ T cells from older (middle-aged) *Tet2^gt^*gene trap hypomorphic mice, which had only 20-30% residual *Tet2* gene expression compared to wildtype CD4^+^ T cells, showed a significant expansion of Tfh cells with hypermethylation of the same intronic region of the *Bcl6* gene (53,54). At first glance, however, these findings contradict the widespread assumption that increased methylation correlates with decreased gene expression. However, a similarly paradoxical finding of increased *BCL6* expression and increased intronic hypermethylation was observed in human B cell lymphoma cells (Raji), which express high levels of *BCL6* compared to plasmacytoma cells (H929) (55) . The authors traced this to decreased binding of CTCF to the hypermethylated region in the first intron of human *BCL6* in Raji cells (55), indicating that the correlation of high BCL6 expression with hypermethylation of the first intron of the mouse *Bcl6* and human *BCL6* genes region is a common feature across species and cell types, whether transformed or untransformed. In fact, mouse CD4^+^ T cells and Treg cells do not express BCL6, and show high CTCF binding in the same region of the first intron of the Bcl6 gene from CTCF ChIP-seq (56) (**Fig. 7A**).

In virally infected wildtype mice, naive CD4^+^ T cells proliferate and differentiate into Th1 and GC-Tfh cells. In contrast, CD4^+^ T cells from *Tet2*-deficient SMARTA TCR transgenic mice (specific for an LCMV GP66–77epitope) showed preferential differentiation into Tfh rather than Th1 cells upon LCMV infection (57,58). FOXO1 and the RUNX1-related transcription factors RUNX2 and RUNX3 are well-known suppressors of Tfh differentiation in mice (48,52), and FOXO1 was shown to coordinate with TET2 at the *Runx2* and *Runx3* loci to mediate their demethylation and expression (58). Genome-wide analysis of Tfh cells from *Tet2*-deficient SMARTA cells in LCMV-infected mice revealed a failure to demethylate the promoter regions of *Runx2* and *Runx3,* resulting in decreased *Runx2* and *Runx3* gene expression (58). These data suggested that TET2 is normally recruited by FOXO1 to *Runx2* and *Runx3* promoters to maintain them in a demethylated and active state. Our RNA-seq data showed that *Foxo1* expression was down- regulated in CD4^+^ Foxp3**^-^** T cells in DKO-severe mice compared to WT (**Suppl. Fig. 1C**), consistent with upregulation of Tfh-related genes; this correlated with increased methylation 3’ of the *Foxo1* transcription start site as determined by 6-base-sequencing (see 5mC tracks in the upper portion of **Fig. 7B**). Thus, unlike the findings for the *Bcl6* gene, increased methylation just 3’ of the *Foxo1* TSS is consistent with decreased expression of the *Foxo1* gene in *Tet2/3 DKO*-severe CD4^+^ YFP(FOXP3)^-^ ex-Treg cells compared to WT naïve CD4^+^ cells (**Suppl. Fig. 1C**). 5hmC in the transcribed region (gene body) of the *Foxo1* gene was high in WT cells (**Fig. 7B**, *middle track*) as shown for essentially all transcribed genes in every cell type examined (43). In our bulk RNA-seq, expression of the *Runx2* and *Runx3* genes were not changed in DKO-severe CD4^+^ YFP(FOXP3)^-^ cells compared to WT (*data not shown*).

TOX2, MAF and BATF are all transcription factors known to promote Tfh differentiation (47,48,59,60), and expression of all these transcription factors was strongly upregulated in DKO-severe mice compared to WT (**Suppl. Fig. 1C, Fig. 2A, Suppl. Fig 6**, *compare RNA-seq tracks*). While the *Tox2* transcription start site (TSS) was largely demethylated in both WT and *Tet2/3 DKO* cells, we observed increased methylation near the TSS and across the *Tox2* gene in *Tet2/3 DKO* compared to WT cells, especially at putative regulatory regions in the 1^st^, 2^nd^ and 3^rd^ introns (**Suppl. Fig. 6A**); this again contradicts the simple hypothesis that increased methylation, especially near putative enhancers or the TSS, is related to decreased gene expression. In contrast, there was no methylation at the 5’ UTR of the *Maf* gene in either WT or DKO cells but decreased methylation in the gene body; the annotation of the *Maf* gene may be incorrect, since transcription appears to start well 5’ of the annotated 5’ UTR (**Suppl. Fig. 6B**). In the case of *Batf*, we observed both increased and decreased methylation at different regions, although RNA expression was unambiguously increased (**Suppl. Fig 6C**). These complex changes are discussed below.

## Discussion

We previously established that loss of FOXP3 expression in Treg cells from *Foxp3-Cre Tet2/3^fl/fl^* mice occurs primarily and progressively in peripheral lymphoid organs, and is driven by loss of TET activity, increased methylation of the conserved intronic *CNS1* and *CNS2* enhancers, and decreased FOXP3 stability (28,33). We show here that FOXP3^+^ Treg cells from these *TET2/3*-deficient mice lose FOXP3 expression and slowly convert to FOXP3^-^ ex-Treg cells with biased (increased) expression of Tfh markers, accompanied by expansion of functional Tfh cells and increased overall inflammation. The markers of increased inflammation included multi-organ lymphocyte infiltration with splenomegaly and lymphadenopathy, a progressive increase in autoantibodies production that was variable with respect to the exact autoreactivity detected, and a progressive increase in the numbers of T cells with an activated phenotype comparing DKO-moderate and DKO-severe *Foxp3-Cre Tet2/3^fl/fl^* mice (**Fig. 1I, J, Fig. 5, Suppl. Figs. 4, 5**). Bulk and single-cell RNA-sequencing of CD4^+^ FOXP3**^-^** T cells from DKO-severe mice showed progressively increased expression of Tfh-associated genes and interferon-stimulated genes (ISGs) in compared to WT mice (**Suppl. Fig. 1C, Fig. 2A**, **Fig. 3C**, **Fig. 6**), a finding confirmed by flow cytometry and immunohistochemistry of the inflamed organs of the *Foxp3-Cre Tet2/3^fl/fl^* mice (**Figs. 4, 5**). Although the correlation between gene expression and DNA methylation changes was complex, we linked Tfh cell expansion and gene expression changes to skewed DNA methylation in TET-deficient FOXP3^+^ Treg cells that converted into FOXP3**^−^**ex-Treg cells with increased expression of BCL6 mRNA and protein, decreased expression of FOXO1, a known suppressor of Tfh differentiation, and a corresponding increase in expression of Tfh markers such as PD-1, CXCR5 and ICOS, (**Fig. 7**, **Suppl. Fig. 6**).

The progressive increase in inflammation observed in DKO-moderate and DKO-severe *Foxp3-Cre Tet2/3^fl/fl^* mice may be explained by our previous finding that Treg cells from these mice lose suppressive activity in a time-dependent manner compared to WT cells (33). This was apparent from the decreased ability of *Tet2/3 DKO* (compared to WT) bone marrow cells to rescue the rampant inflammation and fatal inflammatory phenotype of newborn *scurfy* mice (33). Briefly, functional Tregs do not develop after *scurfy* bone marrow transfer because of a loss-of-function mutation in the *Foxp3* gene (9,61). Hence recipient sub-lethally irradiated *Rag1*-deficient mice administered *scurfy* bone marrow were unable to maintain Treg function, lost weight and died within 50 days; in contrast, co-administered wildtype bone marrow cells prevented autoimmunity completely (33). Co-transfer of *Tet2/3 DKO* bone marrow permitted normal rescue initially, but the mice did not survive past ∼90 days (33). Similarly, transfer of *Tet2/3 DKO* CD4^+^ T cells including Treg cells and ex-Treg cells into normal immunocompetent mice with a full repertoire of functional Treg cells led to increased activation and expansion of bystander T cells in the recipient mice, indicating that the *Tet2/3 DKO* cells conferred a dominant inflammatory phenotype even on immunocompetent mice with normal Treg function (33). This phenomenon resembles the slow onset, but progressive nature, of the autoimmune/ inflammatory phenotype observed in *Tet2/3 DKO* mice. Moreover, the variable yet progressive phenotype of *Tet2/3 DKO* mice resembles that of human patients in whom autoimmune symptoms can manifest variably and may improve and worsen through multiple cycles of remission and relapse.

We also observed a progressive increase of plasma cell frequencies and numbers in *Tet2/3 DKO* mice that was especially apparent in mice with the DKO-severe phenotype (**Fig. 4B, D, F, G**). We attribute this phenomenon to our finding that after loss of FOXP3 (Treg to ex-Treg “conversion”), the resulting “converted ex-Treg” cells expand and acquire the phenotypic and functional features of Tfh cells, including expression of characteristic Tfh genes and surface markers and the functional properties of Tfh cells. The Tfh cells provide help to germinal center B cells, which then differentiate into plasma cells, and plasma cells residing in inflamed tissues produce antibodies in chronic inflammatory and systemic autoimmune diseases. Increased autoantibodies production is indeed observed in *Tet2/3 DKO* mice, especially those categorized as DKO-severe. The parallel increase in cytotoxic CD8^+^ T cells expressing high levels of perforin and granzyme B is also likely due to the emergence of Tfh-like cells that produce IL-21 (**Suppl. Figs 1B, C, Fig. 3C**), which are known to facilitate the differentiate of CD8^+^ T cells into cytotoxic CD8^+^ T cells (62,63). The increased number of neutrophils in spleen and lymph nodes of *Tet2/3 DKO* mice (**Fig. 4B, C)** (33) is less easily explained; it may reflect the known leaky expression of *Foxp3-Cre* expression during hematopoietic development, which occurs at low stochastic levels (64,65). To counter this problem, we are implementing a new strategy for ex-Treg development in which purified Treg cells from *CD4-Cre Tet2/3^fl/fl^*mice (which are not leaky for TET deletion in other lineages) are adoptively transferred into new recipient mice (K. Suzuki, *unpublished*; *to be reported elsewhere*).

Notably, the increased inflammation occurred despite a progressive expansion (increased numbers) of CD4^+^ FOXP3^+^ Treg cells in *Foxp3Cre Tet2/3^fl/fl^* mice (**Fig. 1G**). Heightened expansion in response to ongoing, low-level TCR (or potentially cytokine) stimulation appears to be a characteristic feature of TET-deficient T cells. For instance, we previously reported that expansion of iNKT cells in *CD4-Cre Tet2/3^fl/fl^* mice required antigenic stimulation, since it did not occur in mice lacking the nonclassical MHC protein CD1d, which presents lipid antigens to iNKT cells (34,66). Moreover, bona fide resident Tfh cells show striking expansion in *CD4Cre Tet2/3^fl/fl^* mice (K. Suzuki, *unpublished*; *to be reported elsewhere*).

Changes in 5mC and 5hmC distribution presented a complex picture depending on the genes involved, suggesting that increased methylation due to TET deficiency could reflect direct methylation of essential enhancers or decreased binding of transcriptional repressors such as CTCF. Thus, to understand the influence of DNA methylation changes on gene regulation in depth, it will be necessary to identify the enhancers and transcriptional regulators that govern the expression of each differentially expressed gene, and dissect how DNA methylation at promoters, enhancers and gene bodies influences the binding of these regulators to gene promoters and enhancers. The overall outcome, however, is that CD4^+^ FOXP3^-^ ex-Treg cells from mice with selective deficiency in *Tet2* and *Tet3* in Treg cells show biased conversion to Tfh cells and cause systemic inflammation with variable onset whose severity increases with age.

Based on these data, we postulate the existence of a positive feedback loop that causes a progressive increase in inflammation: when the anti-inflammatory function of Tregs is impaired and Treg proliferation is triggered by some factors such as a minor infection, FOXP3 loss occurs in a cell division-dependent manner, causing inflammation to worsen and prompting the appearance of Tfh cells. This results in further stimulation of CD4^+^ and CD8^+^ T cells through the TCR and the altered cytokine environment (IL2, IL-21, upregulation of *Il2ra* etc) *Tet2/3 DKO* Treg cells proliferate in response to the stimulation and TCR stimulation and proliferation results in a further decrease of FOXP3 stability, promoting the appearance ex-Treg cells with Tfh-like makers and function. This scenario could also hold in humans, suggesting that it would be worthwhile to examine TET activity and regulation in human autoimmune disease.

## Material and Method

### Mice

B6.129(Cg)-*Foxp3tm4(YFP/icre)Ayr/J* (Foxp3-Cre, strain 016959) were obtained from Jackson Laboratory. *Foxp3- Cre Tet2^fl/fl^ Tet3^fl/fl^* mice were generated in our laboratory by crossing *Tet2^fl/fl^ Tet3^fl/fl^* with *Foxp3-Cre* mice. All mice were on the B6 background and maintained in a specific pathogen-free animal facility in the La Jolla Institute for Immunology. Age of the mice used for each experiment was stated in the figure legends. All animal procedures were reviewed and approved by the Institutional Animal Care and Use Committee of the La Jolla Institute for Immunology and were conducted in accordance with institutional guidelines. Euthanasia methods complied with the AVMA Guidelines for the Euthanasia of Animals. Euthanasia was induced by carbon dioxide (CO_2_) asphyxiation/anesthetic overdose coupled with cervical dislocation. The carbon dioxide displacement rate was from 30% - 70% of the chamber volume per minute. For the euthanasia by anesthetic overdose, 5% isoflurane delivered by precision vaporizer. Cervical dislocation was performed as a secondary step to ensure euthanasia.

### Cell preparation and flow cytometry

Single-cell suspensions were prepared from pooled spleen and peripheral lymph nodes (cervical and inguinal) for staining or cell sorting. For analysis of T cell compartments and Treg cell features, single-cell suspensions were stained with anti-mouse antibodies against the following (Clone name, conjugated fluorescence, dilution, manufacturer and catalog number shown in brackets): CD4 (RM4-5, BV421, 1:200, Biolegend, #100548), CD8 (53-6.7, PE-Cy7, 1:200, Biolegend,#100722), TCRβ (H57-597, BV421, 1:200, Biolegend, #109230; H57-597, PE, 1:200, Biolegend, #109208), CD62L (MEL-14, APC, 1:200, Biolegend, #104412), CD44 (PerCP-Cy5.5, 1:200, Biolegend, #103032), B220 (RA3-6B2, APC, 1:200, Biolegend, #103212), Gr1 (RB6-8C5, BV650, 1:200, Biolegend, #108442), CD11b (M1/70, PerCP-Cy5.5, 1:200, Biolegend, #101228), CD98 (RL388, PE-Cy7, 1:200, Biolegend, #128213), CD138 (281-2, BV421, 1:200, Biolegend, #142507), PD-1 (29F.1A12, PE-Cy7, 1:200, Biolegend, #135216; J43, APC, 1:200, Invitrogen, 17-9985-80), CXCR5 (SPRCL5, Biotin, 1:200, Invitrogen, #13-7185-82), BV421 Streptavidin (1:200, Biolegend, #405225) antibodies. For intracellular staining, cells were surface-stained and then stained with anti-IL17A (TC11-18H10.1, PerCP-Cy5.5, 1:100, Biolegend, #506920), anti-IFN-γ (TC11-18H10.1, APC, 1:100, Biolegend, #505810), anti-Perforin (S16009A, PE/Dazzle594, 1:100, Biolegend, #154315), anti-Granzyme B (QA18A28, Pacific Blue, 1:100, Biolegend, #396420), antibodies. The cell were fixed 4% PFA and permed permeabilization buffer (50mM NaCl, 5mM EDTA, 0.02% NaN3, 0.5% TritonX) and analyzed by flow cytometry on Celesta, LSR-II and Fortessa from BD.

### RNA-sequencing library preparation

Total RNA were isolated from CD4^+^ YFP(FOXP3^-^) T cells from pooled spleen and peripheral lymph nodes (cervical and inguinal) from *Foxp3Cre WT* mice and *Foxp3-Cre Tet2/3^fl/fl^* mice (14-weeks-old) using Direct-zol RNA microprep (Zymo, #R2060). RNA-sequencing libraries were prepared using NEBNext Ultra II Directional RNA Library Prep Kit for Illumina (New England Biolabs, #E7760, #7765) according to the manufacture’s protocol and sequenced at the La Jolla Institute sequencing core using Illumina NovaSeq 6000 pair end 150 bp platform.

### RNA-sequencing analysis

Reads were aligned to the mouse reference genome mm10 using STAR version 2.7.11 (1) with the parameters - outFilterMultimapNmax 1 -alignIntronMin 20 -twopassMode Basic -outBAMcompression 10 -alignSJoverhangMin 8 - alignSJDBoverhangMin 1 -outFilterType BySJout -outSJfilterReads Unique -outFilterMismatchNoverReadLmax 0.04 to analyze gene expression. HT-Seq (2) was used to quantify the gene expression levels using the options htseq-count --stranded reverse. Genes with at least ten read counts in one condition were kept for further analysis. Normalization and differential expression analyses were performed using DESEq2 (3). For visualization of the data, we generated tracks using Deeptools94 version 3.5.1, with the option bamCoverage. All related plots were made using R-Studio95 and Integrative Genome Viewer96 version 2.16.0.

### Single-cell RNA-sequencing library preparation and analysis

TCRβ^+^ cells from pooled spleen and peripheral lymph nodes (cervical and inguinal) of *Foxp3Cre WT* and DKO-severe *Foxp3-Cre Tet2/3^fl/fl^* mice (14-weeks-old) were prepared, counted and loaded 10,000 single cells (n=1). Samples were processed per the manufacturer’s protocol using Chromium Next GEM Single Cell 3’ Kit v3.1 (10X Genimics) and sequenced were conducted at the La Jolla Institute sequencing core on an Illumina Novaseq 6000 sequencer using a 100-bp kit. Samples were sequenced using a 28|10|10|90 run configuration. Sample demultiplexing, barcode processing, alignment, filtering, UMI counting and aggregation of sequencing runs were performed using the Cell Ranger analysis pipeline (v.7.1.0) and the mouse reference genome mm10. Downstream analysis was performed using the R package Seurat (4).

### 6-base sequencing (simultaneous detection of 5hmC and 5mC by duet multiomics solution evoC (biomodal)) library preparation and analysis

Naïve CD4^+^ T cells from *Foxp3Cre WT* mice (14-weeks-old) and Tfh cells from DKO-severe *Foxp3-Cre Tet2/3^fl/fl^* mice (14-weeks-old) were sorted and extracted genomic DNA using DNeasy Blood & Tissue Kit (Qiagen, #69504). The duet multiomics solution evoC method (6-base sequencing) was performed according to the manufacturer’s instructions (duet exoC User guide v.3), using 60-70 ng of sonicated genomic DNA as starting material (n=2) Data were processed as in ref. (5) and the mean coverage for the samples ranged from 10x to 20x. Libraries were sequenced at the UCSD Institute for Genomic Medicine Genomics Center using Illumina NovaSeq X Plus pair end 150 bp platform.

### Histology

Pancreas, colon, skeletal muscle, heart, liver, lung, salivary gland, brain, knee joint, stomach were isolated from 14- week-old *Foxp3Cre WT* mice and *Foxp3-Cre Tet2/3^fl/fl^* mice. Samples were fixed using 10% zinc formalin, washed in 70% isopropanol, processed and embedded in paraffin. The blocks were then cut and 4 µm sections were stained with haematoxylin and eosin (H&E) (6). Slides were digitized with ZEISS AxioScan.Z1 scanner equipped with a 20x 0.8NA objective. Whole slide images were analyzed by a board-certified veterinary pathologist in QuPath (7) software. Neoplastic changes were not seen and inflammation was scored on a 0-5 scale for quantity of infiltrating cells including neutrophils, lymphocytes, macrophages, and plasma cells in specified regions of the analyzed tissues.

### Orion multiplexing

Five-micron sections were placed on positively charged slides, dried, and baked at 60°C for 1h to adhere tissue, then deparaffinized using three cycles of Pro-Par clearant dipping (20×) and submersion (10min). Slides underwent rehydration in reagent alcohol: twice in 100% (20× dips, 1.5min submersion) and once in 90% (20× dips, 1.5min submersion), followed by a 2min DI water rinse. Antigen retrieval was performed in pH 6 citrate buffer in a Biocare Medical Decloaking Chamber™ NxGen (Program 5: 110°C, 15min) and then cooled to RT. Autofluorescence was quenched in PBS with 2.4 mM NaOH and 1.47 M H_2_O_2_ solution under LED (1h) and UV light (30min). Slides were placed in a Freequenza rack (8), blocked with Image-iT FX Signal Enhancer (15min), and washed (1mL, 0.025% TritonX-100 in PBS). Slides were stained with a 16-plex conjugated antibody panel, including anti-mouse CD3ε (D4V8L)-ArgoFluor 548, CD8α (4SM15)-ArgoFluor 658, B220 (RA3-6B2)-ArgoFluor 698, IRF4 (E8H3S)-ArgoFluor 706, FOXP3 (D6O8R)-DIG-ArgoFluor 760, CD4 (4SM95)-ArgoFluor 812, CD31 (D8V9E)-ArgoFluor 874 and others in Candor Antibody Stabilizer PBS with 5% mouse and 5% rabbit serum overnight at 4°C. After warming to RT and washing (4mL), slides were stained with secondary antibodies in Antibody Stabilizer PBS with 10% goat serum (30min) and washed (4mL). Slides were counterstained with Hoechst (1:1000, 5min), washed (5mL PBS), coverslipped with ArgoFluor Mounting Medium, and dried for 48h before scanning. Images were acquired with a 20× 0.75 dry objective (0.325μm/pixel) using whole-tissue tiling. The Orion system (RareCyte, Seattle, Washington, USA) captured ∼10nm emission bands via angled dichroic mirrors. Processing used Rarecyte’s algorithm for tile stitching, non-linear channel alignment, and spectral unmixing based on previously acquired single-color controls. The Orion images were analyzed in QuPath 0.6.0 (9). For cell counting, images were analyzed in QuPath 0.6.0 (9). Briefly, cells were segmented using InstanSeg (10). Random forest machine learning classifiers were trained to identify cells positive for each of ICOS, CD3 (hi vs low), CD4, CD8, and IRF, as well as cells simultaneously negative for CD3ε, B220, and F4/80. Ex-Tregs were defined as ICOS^+^ CD3ε^+^ CD4^+^ FOXP3^-^ cells. Plasma cells were defined as IRF^+^ CD3ε^-^ B220^-^ F4/80^-^.

### Measuring auto-antibody level in serum

Blood was collected by cardiac puncture in BD Microtainer (BD, #365978) and serum was isolated. Autoantibodies level in serum was measured using Human Autoimmune Disease IgG Autoantibody array G1 (RayBiotech, #PAH- AIDG-G1-16) according to the manufacture’s protocol. We used biotin-conjugated Anti-mouse IgG (Jackson ImmunoResearch, #115-065-071) instead of biotin-conjugated Anti-human IgG to measure mouse auto-antibodies.

### Quantitative real-time PCR

Total RNA was isolated using Direct-zol RNA microprep (Zymo, #R2060); cDNA was synthesized using SuperScript™ IV First-Strand Synthesis System (Thermo Fisher, #18091050). Quantitative real-time PCR was performed using PowerUp™ SYBR™ Green Master Mix for qPCR (Thermo Fisher, #A25742) on a StepOnePlus real-time PCR machine (Applied Biosystems). Gene expression was normalized to *Gapdh*. Primers used to detect the expression levels of *Tet2* are as following:

Tet2 forward primer: AACCTGGCTACTGTCATTGCTCCA

Tet2 reverse primer: ATGTTCTGCTGGTCTCTGTGGGAA

*Gapdh* forward primer:TCACCACCATGGAGAAGGC

*Gapdh* reverse primer: GCTAAGCAGTTGGTGGTGCA

### Statistics

*P* values from one-way ANOVA test were used for all the statistical comparisons between different groups and data were displayed as mean ± SEM. (Prism). *P* values are denoted in corresponding figures as: \**P* < 0.05, \*\**P* <0.01, \*\*\**P* < 0.001, \*\*\*\**P* < 0.0001.

## Figure legends

**Figure S1.**
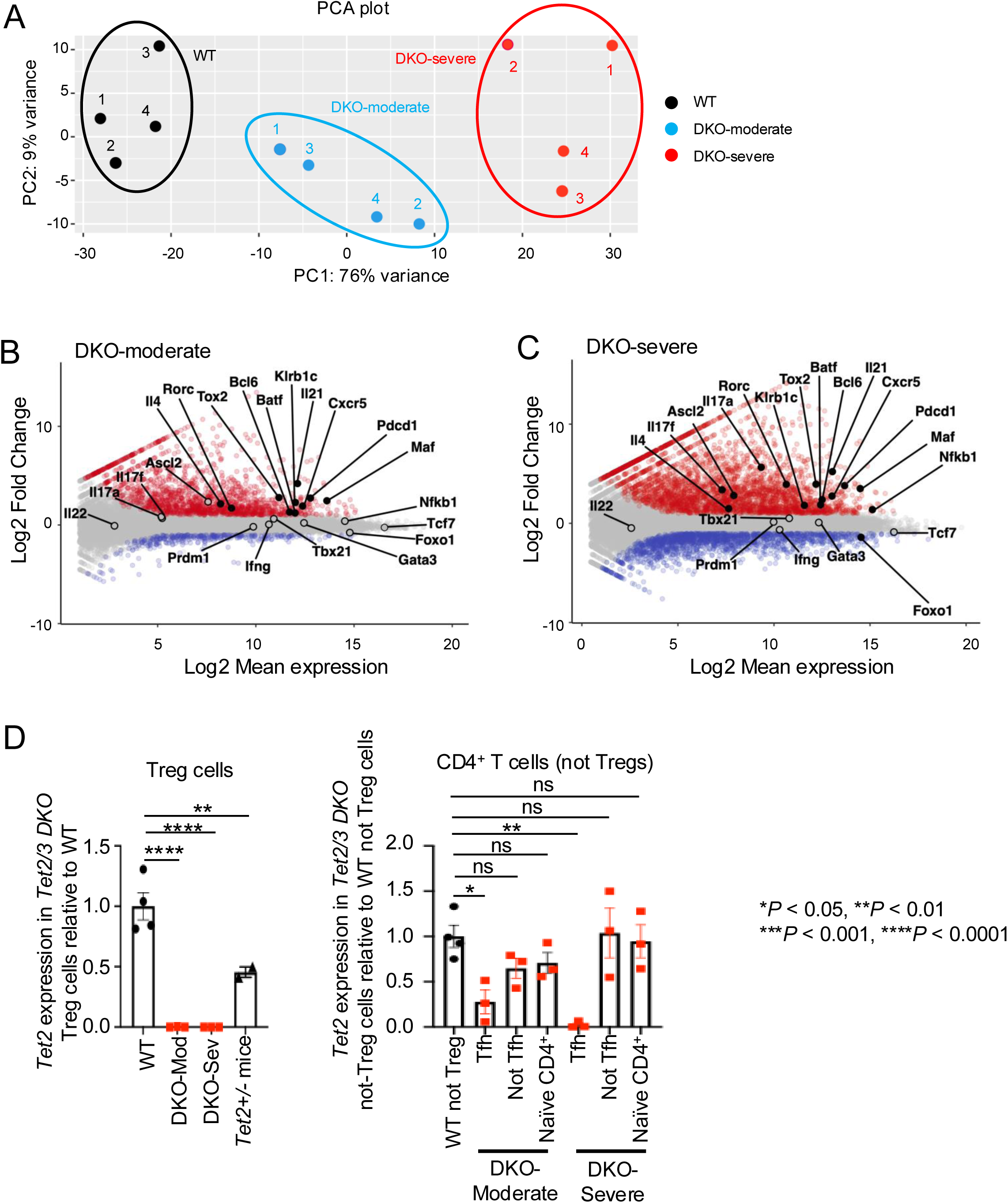
**A-C.** RNA-seq analysis for CD4^+^ YFP(FOXP3^-^) T cells from *Foxp3Cre WT* mice and *Foxp3-Cre Tet2/3^fl/fl^* mice (14- weeks-old). **A.** PCA plot. **B-C.** Mean average (MA) plot of genes differentially expressed in DKO-moderate relative to their expression in WT (**B**), DKO-severe relative to their expression in WT (**C**). **D.** The graphs show the *Tet2* expression in *Tet2/3 DKO* Treg cells relative to WT Treg cells (*left*), the *Tet2* expression in *Tet2/3 DKO* not-Treg cells relative to WT not Treg cells (*right*). Treg cells: CD4^+^ YFP(FOXP3)^+^, WT not Treg cells: CD4^+^ YFP(FOXP3)^-^, Tfh cells: CD4^+^ YFP(FOXP3)^-^ PD-1^+^ CXCR5^+^, not Tfh cells: CD4^+^ YFP(FOXP3)^-^ PD-1^-^ CXCR5^-^, Naïve CD4^+^ cells: CD4^+^ YFP(FOXP3)^-^ CD62L^high^ CD44^low^.

**Figure S2.**
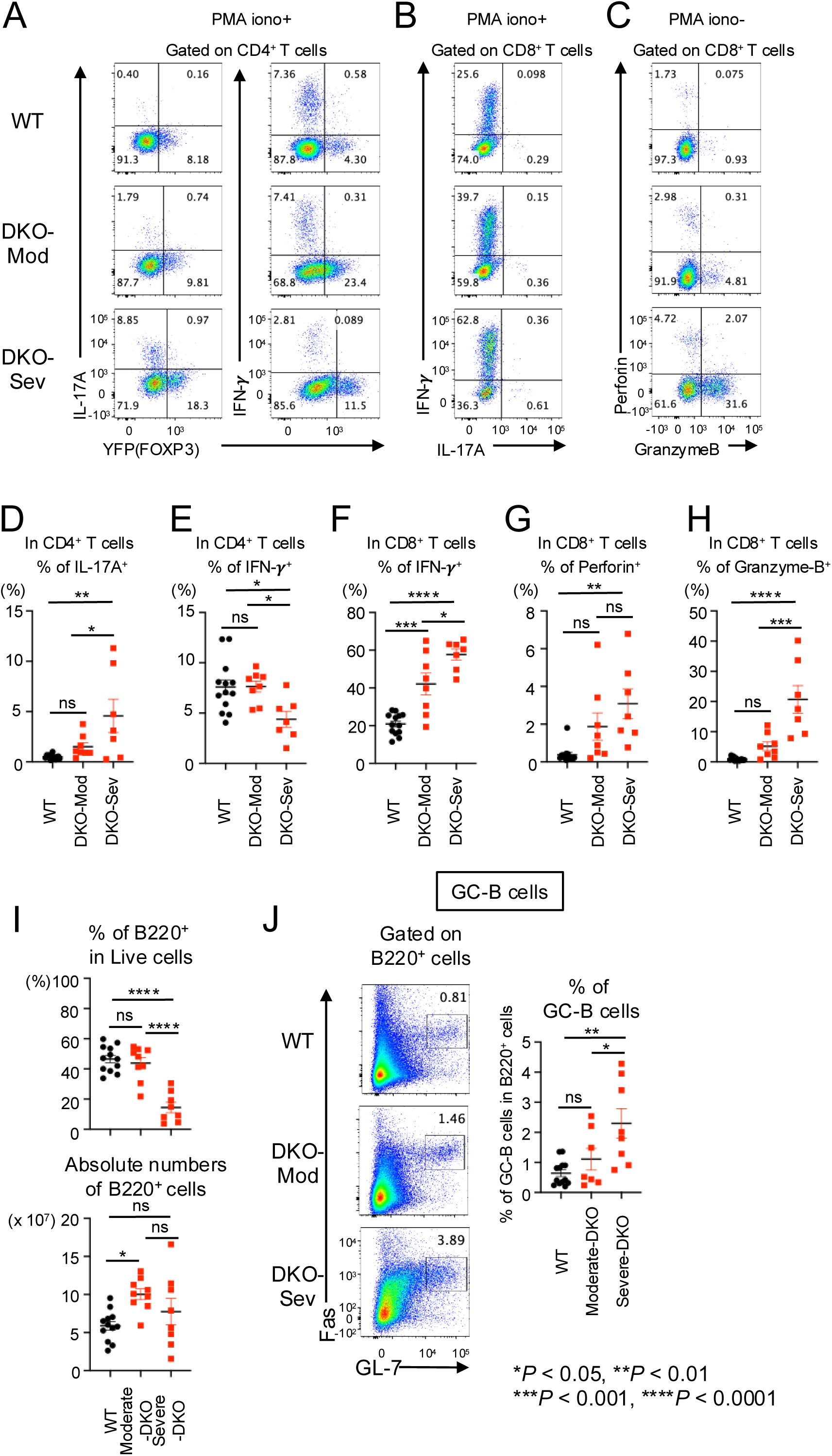
**A-C.** Flow cytometry analysis of cytokines in CD4^+^ and CD8^+^ cells. IL-17A^+^ cells and IFN-γ^+^ cells with PMA and ionomycin stimulation (gated on TCRβ^+^ CD4^+^ cells) (**A**), IL-17A^+^ and IFN-γ^+^ cells with PMA and ionomycin stimulation (gated on TCRβ^+^ CD8^+^ cells) (**B**), Perforin^+^ and Granzyme B^+^ cells (gated on TCRβ^+^ CD8^+^ cells) (**C**) in pooled spleen and peripheral lymph nodes (cervical and inguinal) of from 14-weeks-old *Foxp3Cre WT* and *Foxp3-Cre Tet2/3^fl/fl^* mice. **D-E.** Quantification of the frequency of IL-17A^+^ cells (**D**), IFN-γ^+^ cells (**E**) in CD4^+^ T cells. **F-H.** IFN-γ^+^ (**F**), Perforin^+^ cells (**G**), Granzyme B^+^ cells (**H**) in CD8^+^ T cells. **I.** Quantification of the frequency of B220^+^ cells in live cells and absolute number. **J.** Flow cytometry analysis of GC-B cells (Fas^+^ GL-7^+^ cells) (gated on B220^+^ cells) in pooled spleen and peripheral LNs (cervical and inguinal) from 14-weeks-old *Foxp3Cre WT* and *Foxp3-Cre Tet2/3^fl/fl^*mice (*left*). Quantification of the frequency of Fas^+^ GL-7^+^ cells in B220^+^ T cells.

**Figure S3.**
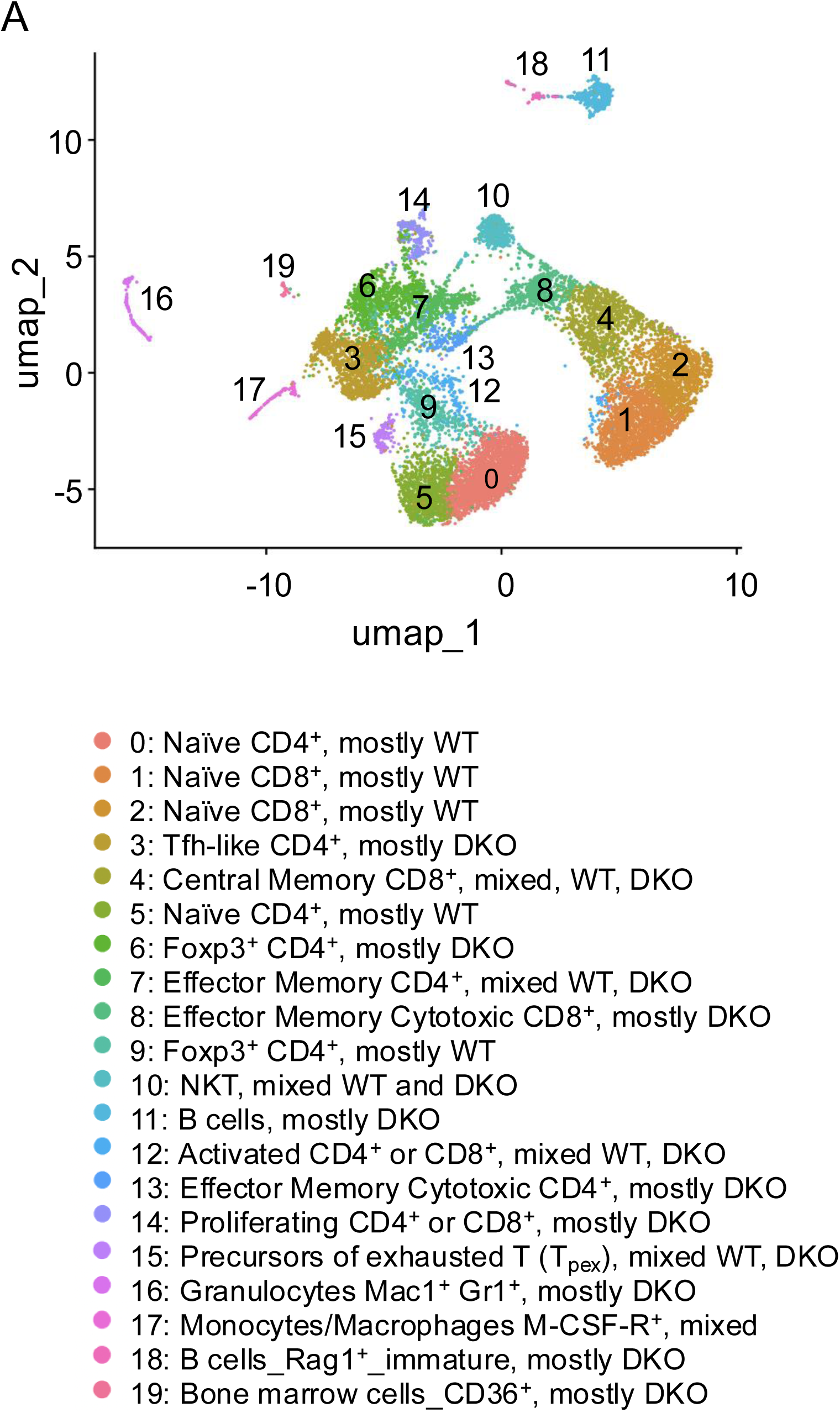
**A.** The UMAP plot showed 0-19 clusters from sc-RNA sequencing. Annotated clusters are shown. TCRβ^+^ cells were sorted from pooled spleen and peripheral lymph nodes (cervical and inguinal) from *Foxp3Cre WT* mice and *DKO- severe Foxp3-Cre Tet2/3^fl/fl^* mice (14-weeks-old) and sc-RNA sequencing was performed (n=1).

**Figure S4.**
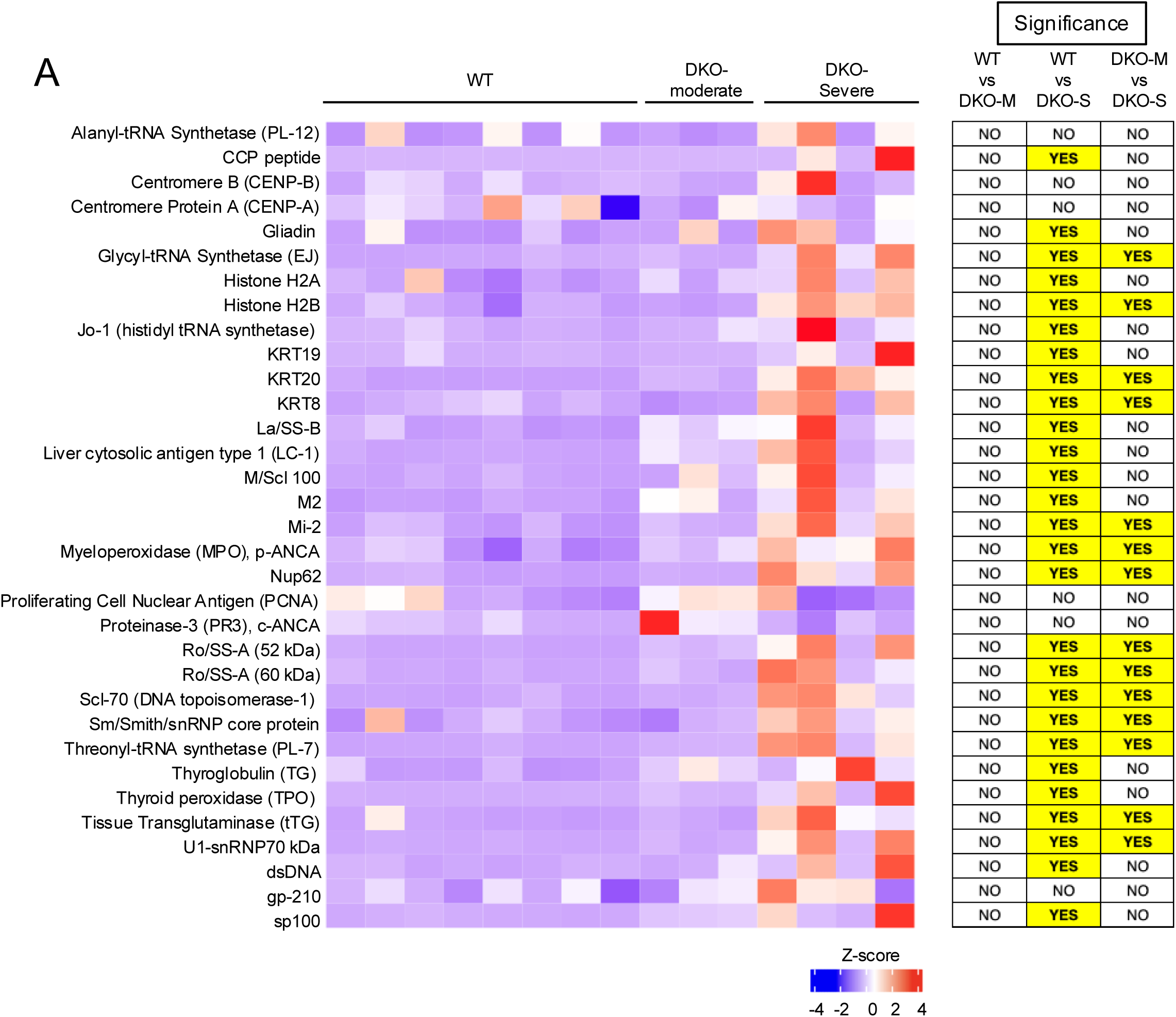
**A.** Heatmap from polyreactivity assay for serum isolated from WT (n=8) and DKO-moderate (n=3), DKO-severe (n=4) tested by autoantibody array assay kit (13-15 weeks old mice). For data analysis, R package ‘limma’ and multiple comparisons correction was performed (adjusted p value < 0.05 between tested groups).

**Figure S5.**
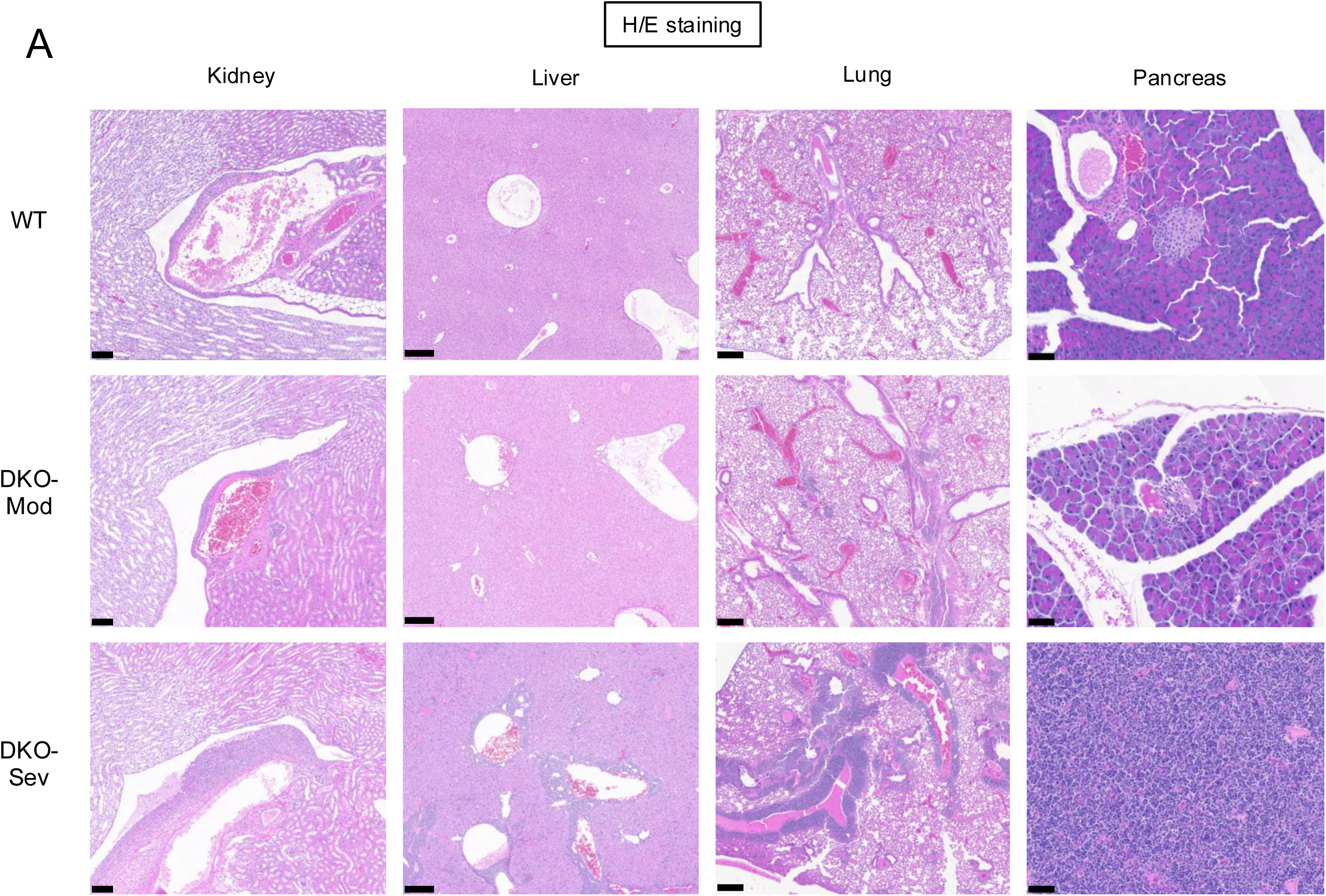
**A.** H&E staining of Kidney, Liver, Lung and Pancreas from 14-week-old *Foxp3Cre WT* and *Foxp3-Cre Tet2/3^fl/fl^* mice. Scale bar; Kidney: 100μm, Liver: 200μm, Lung: 250μm, Pancreas: 50μm.

**Figure S6.**
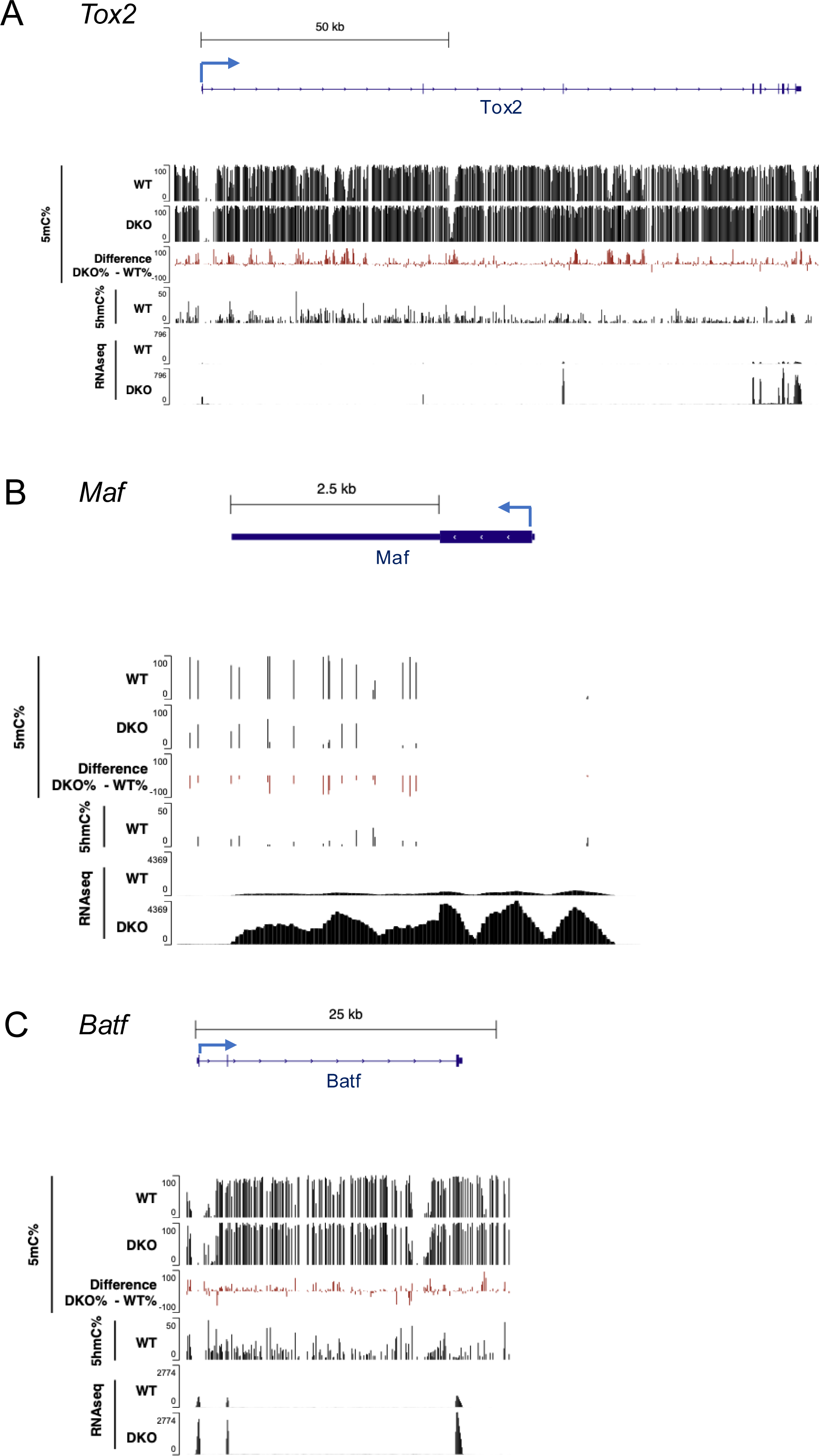
**A-C.** Genome browser views showing 5mC% (Track 1), 5hmC% (Track 2) from 6 base sequencing, gene expression (RNA-seq, track 3) in *Tox2* (**A**), *Maf* (**B**) and *Batf* (**C**) locus. 6 base sequencing; WT: naïve CD4^+^ T cells (CD4^+^ YFP(FOXP3)^-^ CD62L^high^ CD44^low^) from *Foxp3Cre WT* mice, DKO: Tfh like cells (CD4^+^ YFP(FOXP3)^-^ PD-1^+^ CXCR5^+^) from DKO-severe *Foxp3-Cre Tet2/3^fl/fl^* mice. RNA seq; WT: CD4^+^ YFP(FOXP3)^-^ T cells from *Foxp3Cre WT* mice, DKO: CD4^+^ YFP(FOXP3)^-^ T cells from DKO-severe *Foxp3-Cre Tet2/3^fl/fl^* mice.

## Acknowledgement

We thank Q. Li (University of Pennsylvania) for sharing his curated list of ISGs. We also thank the Department of Laboratory Animal Care and the animal facility for excellent support, and C. Kim, S. Alarcon, K. Kim, Z. Mikulski, S. McArdle, S. Goldstein and colleagues of the La Jolla Institute Flow Cytometry, Next-Generation Sequencing *(RRID:SCR_023107),* Histopathology and Microscopy Core Facilities for help with cell sorting, next-generation sequencing, and microscopy, and immunostaining, respectively. The FACSAria II Cell Sorter (S10RR027366), the NovaSeq 6000 (S10OD025052), the Zeiss LSM 880 (S10OD030417) and LSM 980 (S10OD030417) were acquired through the Shared Instrumentation Grant Program. This work was funded by NIH grants R01 AI128589 and R35 CA210043 to A.R., NIAID AI109842, AI040127 to P.G.H., the Tullie and Rickey Families SPARK Awards for Innovations in Immunology at La Jolla Institute to K.S. and I.F.L-M, and an Astellas Foundation Award for Research on Metabolic Disorders, a Mochida Memorial Foundation Award for Medical and Pharmaceutical Research, and KAKEN grant JP23K24147 from the Ministry of Education, Culture, Sports, Science and Technology (MEXT Japan) to A.O. H.S. was supported by the Pew Latin-American Fellows Program from The Pew Charitable Trusts, by a Fellowship from the California Institute for Regenerative Medicine and a PEW Repatriation Award.

## Author contribution

K.S. performed all the experiments (flow cytometry, library preparation, autoantibody array assay etc.) and assembled the figures. L.J.A.-V., B.V.R., L.H.-E. and I.F.L.-M performed bioinformatics analyses (RNA-seq, sc- RNA-seq, 6-base seq). H.S. and P.G.H. supervised the bioinformatic analyses by L.J.A.-V. and B.V.R. A.O. annotated the clusters in sc-RNA seq. A.L.-C. analyzed genome compartmentalization using data from a Hi-C experiment on induced Treg cells supervised and analyzed by F.A. and performed by D.S.-C. A.R. directed the experiments and data analyses. K.S. and A.R. wrote the paper with input from all authors.

